# Divergent condensates tune transcriptional responses during stress

**DOI:** 10.64898/2026.02.12.705659

**Authors:** Jeffrey N. Dudley, Joel E. Berends, Chayan K. De, Tongchen He, Giovana B. Veronezi, Benedict Abdon, Anjali Sengar, Matthias C. Truttmann, Mats Ljungman, Lanbo Xiao, Srinivas Ramachandran, Sethuramasundaram Pitchiaya

## Abstract

Dynamic reorganization of the transcription machinery within nuclear membrane-less compartments is an emergent feature of mammalian stress response, associated with critical cellular decisions. However, mechanisms governing the subcellular formation of these stress-induced condensates and their role in transcription regulation remain poorly understood. Here, we find that heat shock factor 1 (HSF1), transcriptional mediator of protein and cellular homeostasis, forms condensates during various adverse conditions, but these assemblies exhibit context-dependent divergent transcriptional outcomes. During heat shock, HSF1 orchestrates the coordinated assembly of transcription hubs via canonical activation, including post-translational modifications (PTMs), trimerization, and DNA binding. While HSF1’s disordered regions restrict condensate formation in unstressed situations, they promote stress-induced condensate maturation to transcriptionally active states. Strikingly, HSF1 condensates that form during other environmental and chemotherapeutic stresses stall at distinct stages of hub formation, assemble independent of PTMs, and exhibit reduced sub-condensate dynamics. These aspects culminate in attenuated genomic occupancy and transcriptional output at HSF1-associated loci, consistent with functional impairment of HSF1 and the transcription machinery via sequestration. Our work suggests that stress-induced transcription factor condensates drive conserved responses during physiological perturbations, but can be inactivated during pathological insults, rationalizing HSF1 and transcriptional dysfunction across degenerative diseases and toxic exposures.

## INTRODUCTION

Membrane-less compartments drive numerous cellular functions^1^. Specifically, disruption of cellular homeostasis results in the rapid reorganization of biomolecules within numerous such subcellular condensates in all forms of life^2–4^. These stress-induced condensates mediate various emergent processes, including stress sensing, and are associated with crucial cell fate decisions, especially cell survival during adverse conditions^2–6^. However, aberrant condensation is linked to numerous degenerative diseases in humans^7,8^, underscoring condensates’ importance in physiological responses and pathology. In eukaryotes, accumulation of transcription factors (TFs) and the transcriptional machinery within nuclear condensates are prominent, conserved features of stress^9–11^. While these condensates are posited to facilitate transcription regulation^10,11^, their precise role in transcriptional response to stress remains unclear.

Heat shock factor 1 (HSF1) is an evolutionarily conserved, archetypical stress-induced TF that is central to protein and cellular homeostasis^12,13^. In response to a multitude of environmental stresses^13–20^, including heat shock (HS)^13,21^, HSF1 undergoes extensive post-translational modifications (PTMs), trimerizes and binds to specific genomic elements termed heat shock elements (HSEs)^12^. This activated HSF1 drives its eponymous gene regulatory program called the heat shock response (HSR), which involves chaperone gene induction among other things, to counteract stress-induced proteotoxicity^12,13^ and beyond. Dampened HSF1 levels are associated with chronic diseases, infertility and neurodegenerative diseases^12,22,23^, and overexpression or overactivity of HSF1 is a feature of cancer^12^, wherein malignant cells exhibit non-oncogene addiction to HSF1^24^. While the role of HSF1 and the HSR in adaptive stress response is well defined in yeast^25^, human HSF1 drives context-(stress- and cell-)specific regulatory programs^13,14,18^, highlighting our incomplete understanding of HSF1’s vast functions in stressed human cells. Adding to its functional complexity, HSF1 drives cancer specific regulatory programs that are distinct from the HSR^26^, indicating distinct roles in pathology.

HSF1 accumulates within nuclear condensates during a variety of stresses^9,17,18,20,21^ and specifically in tumor cells of patient tissues^27^. Based on studies in yeast^28^ and in HS-subjected mammalian cells, manifestations of HSF1 as mesoscale nuclear stress bodies (nSBs)^29^ and phase separation of HSF1 within nanoclusters^21^ are predicted to represent transcriptional condensates. Yet mechanisms driving the formation and transcriptional function of these HS-induced HSF1 condensates remain incompletely understood in human cells. Moreover, HSF1 has been reported to undergo phase transition *in vitro* during oxidative stress^17^ and in cancer cells upon proteotoxic drug treatment^27^, suggesting that HSF1 condensates may exhibit variable transcriptional activity across distinct physiological and pathological scenarios in human cells, but this thesis remains to be tested. Unlike cytoplasmic stress granules that sequester translationally repressed mRNAs in all settings^7^, it is unknown if stress-induced HSF1 condensates have a consensus function in transcription.

Here, we use an array of genetic and chemical perturbations, quantitative high-resolution microscopy, single-molecule tracking, epigenomics and nascent RNA sequencing to functionally dissect HSF1 condensates during a variety of physiologically and pathologically relevant perturbations. Specifically, stresses like acute HS result in transcriptionally hyperactive HSF1 condensates, whereas several other insults accumulate HSF1 within transcriptionally impaired condensates. HS-induced HSF1 condensates bear hallmarks of transcription hubs, enriching for co-activators of (super-)enhancers^30^. However, condensates that form during other adverse environmental and chemotherapeutic stresses do not contain the full complement of factors required for active hubs. At the intramolecular level, DNA binding and trimerization of HSF1 are necessary for condensate formation across all stresses. Strikingly, the predominantly disordered regulatory domain prevents condensate formation in the absence of stress. However, the regulatory domain along with the disordered activating domain recruit distinct factors that constitute active hubs during stress, essentially mediating condensates’ transcriptional function.

Moreover, HS-induced active hubs require activating PTMs for their formation, but non-HS stress-induced condensates are PTM-agnostic. Consistent with transcriptional differences, intra-condensate HSF1 mobility was significantly higher in HS-induced condensates, as expected for dynamic responses, compared to other stresses, wherein relatively immobile HSF1 resides in stalled hubs. Finally, we find that condensate activity tightly correlates with genome-wide HSF1 chromatin binding and nascent transcriptomic output, highlighting elevated transcriptional responses during HS and relatively muted responses in other stresses. Overall, we find that nuclear HSF1 condensation is a hallmark of stress, but these condensates exhibit stress-type-dependent divergent functionality. Our work suggests that stress-induced TFs form dynamic transcriptional condensates to mount robust responses during physiological perturbations. However, the TF themselves are vulnerable during various pathological insults, resulting in their suppressed function via sequestration. These findings rationalize functional inactivation sans downregulation of stress-responsive TFs as an additional driver of several idiopathic degenerative diseases and maladies driven by environmental toxins.

## RESULTS

### Divergent roles of HSF1 condensates

HSF1 has been reported to form nuclear foci in mammalian cells during a variety of acute stresses^9,17,20,21,27^. To dissect cellular features of these condensates, we used high-resolution imaging of endogenous human HSF1 using immunofluorescence (IF) and 3-dimensional highly inclined laminated optical sheet microscopy (3D-HILO, Methods) in U2OS (osteosarcoma) cells that have been treated with various acute insults (Fig. 1a-b and Supplementary Fig. 1a-j). Specifically, we subjected cells to various types of stressors that have been shown to induce HSF1 foci in mammalian cells, namely heat shock (HS, acute proteotoxic stress model^9,21^), azetidine carboxylic acid (ACA, plant amino acid analog toxin that drives proteotoxicity in humans^9^), cadmium chloride (CdC, heavy metal poison^9,20^), sodium arsenite (SA, metalloid toxin and oxidative stress model^18^) and hydrogen peroxide (HP, oxidative stress model^17^). Based on the induction of chaperones^14–16,19^, we posited that a few other environmental, metabolic and chemotherapeutic stressors that are typically encountered by human cells will also induce HSF1 condensates. Thus, we additionally subjected cells to cobalt chloride (CC, heavy metal poison and chemical hypoxia model^31^), methylglyoxal (MGO, disease associated metabolite and model glycating agent^15^) and arsenic trioxide (ATO, chemotherapeutic agent^19^). By adapting deep-learning based segmentation models (Supplementary Fig. 1a, Methods) we extracted morphometrics (Supplementary Fig. 1b-f) of HSF1 condensates. We find that nuclear HSF1 is relatively diffuse under unstressed conditions but reorganize within nano-to-mesoscale condensates in all stressful scenarios (Fig. 1a-b), irrespective of antibodies used to probe HSF1 (Supplementary Fig. 1g-j), and this aspect is preserved in *C. elegans* (Supplementary Fig. 1k-m). However, the extent of condensate formation and morphology of condensates were significantly variable across stresses (Fig. 1a-b), indicative of functional differences.

**Figure 1.**
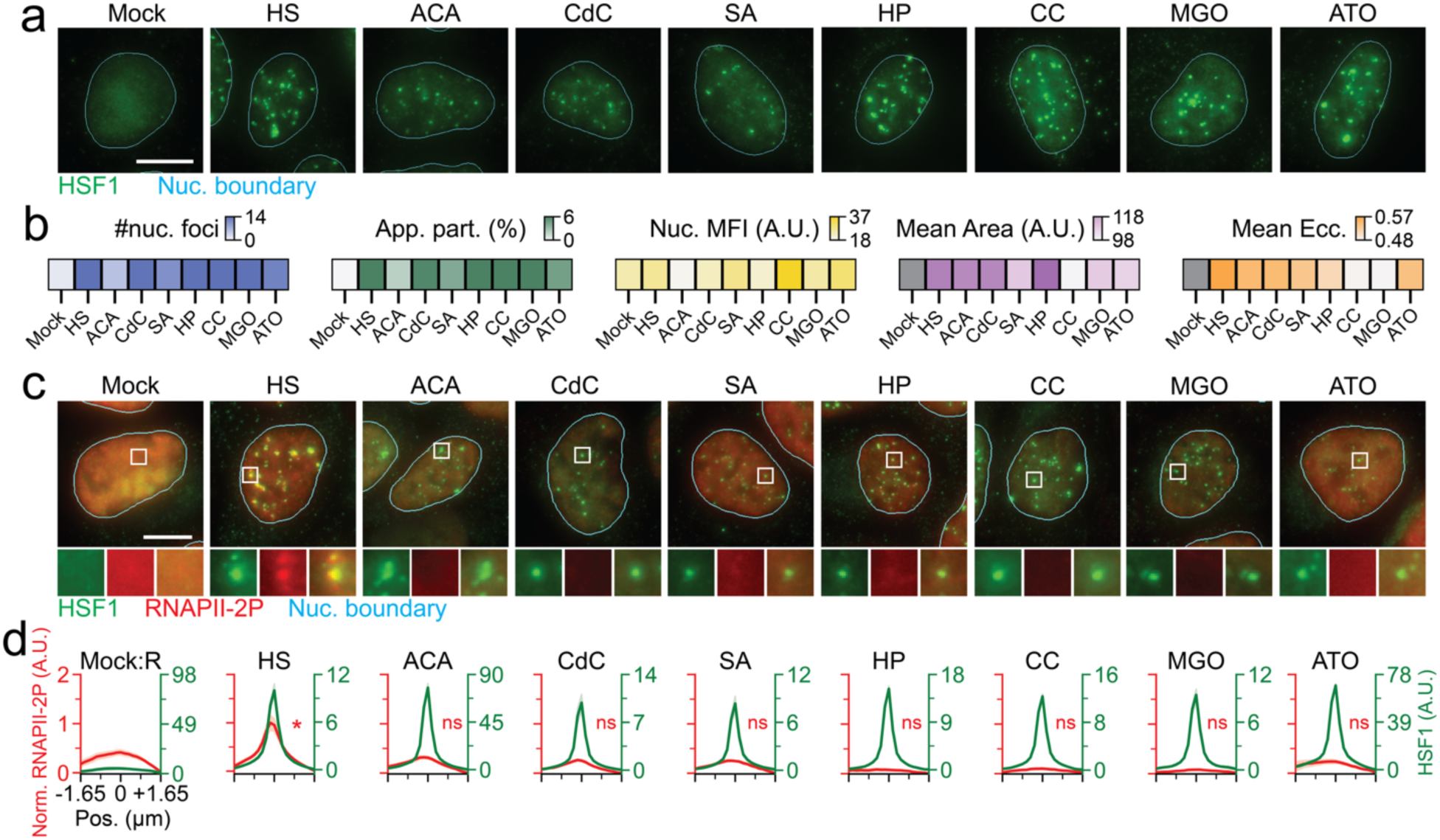
Nuclear HSF1 condensates, a hallmark of cellular stress, exhibit context-dependent divergent transcriptional competency. (a) Representative pseudo-colored IF images of U2OS cells Mock treated (37 °C) or stressed (HS, 42 °C, 1 h; ACA, 10 mM, 6 h; CdC, 1 mM, 4 h; SA, 0.5 mM, 1 h; HP, 2 mM, 1 h; CC, 10 mM, 4 h; MGO 10 mM, 1 h; ATO 0.5 mM, 1 h) stained for HSF1 (green). Nuclear (Nuc.) boundary (cyan) obtained from segmentation of DAPI signal. Scale bar, 10 μm. (b) Heatmaps of mean number of HSF1 foci, HSF1 apparent partition (App. Part.), Nuc. HSF1 mean fluorescent intensity (Nuc. MFI), mean HSF1 foci area, and mean HSF1 foci eccentricity (Ecc.) per nucleus from (a). Color scale for each is also represented (n ≥ 73 cells, 1138 condensates, per condition). (c) Representative pseudo-colored IF images of U2OS cells treated with conditions from (a) and stained for HSF1 (green) and RNAPII-2P (red). Nuc. boundary (cyan) obtained from segmentation of DAPI signal. Scale bar, 10 μm. Zoom-ins are 2.7 μm × 2.7 μm. (d) Line scan metaplots of RNAPII-2P signal across segmented HSF1 condensates for each condition in (c). *=p<0.05, ns = not significant by two-sided unpaired T-test, (n ≥ 78 cells, 1958 condensates, per condition).

We then used co-IF and 3D-HILO to image endogenous HSF1 and RNA polymerase II that is phosphorylated at the 2^nd^ serine of its C-terminal domain repeat (CTD, a marker of actively transcribing RNAPII^32^, i.e. RNAPII-2P), to visualize transcription competency of HSF1 condensates across stresses. Meta-analysis across thousands of condensates in stressed cells and comparative random nuclear loci in unstressed cells (Supplementary Fig. 2a-b) showed that RNAPII-2P was enriched at HS-induced HSF1 condensates, but not in any of the other stresses (Fig. 1c-d). As expected for stress-induced global transcription reduction, RNAPII-2P signal was diminished across all stresses (Supplementary Fig. 2c), suggesting that RNAPII-2P concentration is not a limiting factor for differential co-enrichment in HSF1 condensates across stresses. Concordantly, analysis of HSF1 condensates with similar partitioning (Supplementary Fig. 2d-e) preserved differences in transcription competency between HS and other stresses. These results suggest that HSF1 condensate formation is an evolutionarily conserved feature of stressful exposures, but condensates are transcriptionally competent only during certain contexts. Based on these observations, we categorize stresses and the appropriate condensates formed in these distinct contexts as Type I (transcriptionally competent) and Type II (transcriptionally sub-/incompetent).

### HSF1-driven transcription hubs in HS

Considering that HSF1 condensates exhibit robust transcriptional capacity during acute HS, we first sought to dissect the mechanism of transcription activation in this condition. Using co-IF coupled with 3D-HILO, we found that HS-induced endogenous HSF1 condensates do not enrich for H3K4me3, a canonical histone modification at gene promoters^33^ (Fig. 2a-b). However, these condensates were enriched for factors that drive enhanced transcription, namely H3K27ac, a histone modification associated with elevated transcription and enriched at enhancers^33^, BRD4, factor that binds acetylated histones^34^, and MED12, component of the mediator complex that connects distal chromatin elements^35^ (Fig. 2a-b). As expected, RNAPII and RNAPII-5P, a post-translationally modified version that indicates transcription initiation^32^, were also enriched at HSF1 condensates (Fig. 2a-b). We then tested if HSF1 is required for the formation of such transcription competent condensates. To this end, we generated U2OS cells that lacked HSF1 using CRISPR-Cas9 mediated gene editing (UHKO, Supplementary Fig. 3a-b) and subjected them to HS. Using high-resolution imaging at high-throughput (HRIHT) that images 50-100 cells per field-of-view (at ∼81 nm resolution) and IF for endogenous content, we found that H3K27ac, BRD4, MED12 and RNAPII formed mesoscale foci during HS in U2OS cells (Fig. 2c-d). Strikingly, signal from all these components were completely dispersed during HS in UHKO cells (Fig. 2c-d). This trend was consistent in multiple cell lineages (U2-OS, MCF7 and HeLa) that were depleted for HSF1 by silencing RNAs (Fig. 2e-f and Supplementary Fig. 3c-k). Using a doxycycline (dox) controlled stably transduced cell system, we re-expressed full-length (FL) Halo-tagged HSF1 in UHKO cells (UHKO-HaH) to levels that matched endogenous content found in U2OS cells (Supplementary Fig. 3l-m). We found that Halo-HSF1 formed condensates during HS (Supplementary Fig. 3n) and mediated the re-organization of transcription machinery within condensates (Fig. 2g-h). These data strongly support the notion that HSF1 is required for the formation of transcription hubs^36^ during HS.

**Figure 2.**
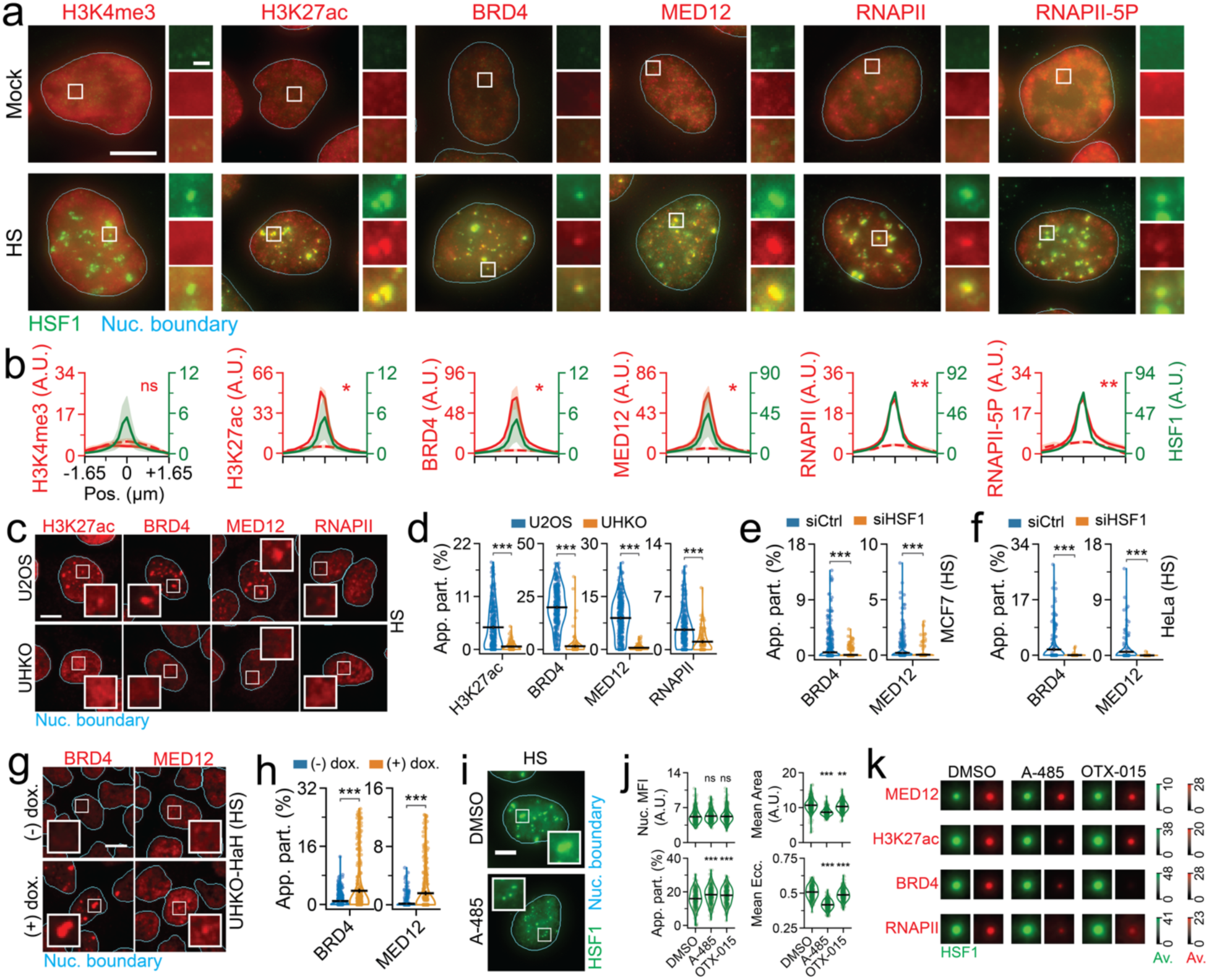
HSF1 drives transcription hub formation during HS. (a) Representative pseudo-colored IF images of Mock treated or HS subjected U2OS cells and stained for HSF1 (green) and H3K4me3, H3K27ac, BRD4, MED12, RNAPII, or RNAPII-5P (red). Scale bar, 10 μm. Zoom-ins are 2.7 μm × 2.7 μm with 1 μm scale bar. (b) Line scan metaplots of co-IF signal across segmented HSF1 condensates from (a). Dashed red lines represent co-IF signal from spatial randomizations. *=p<0.05, **=p<0.005, ns = not significant, by two-sided unpaired T-test (n ≥ 53 cells, 1750 condensates, per condition). (c) Representative pseudo-colored IF images of U2OS or UHKO cells subjected to HS and stained for H3K27ac, BRD4, MED12, or RNAPII (red). Scale bar, 10 μm. Zoom-ins are 5 μm × 5 μm. (d) Quantification of H3K27ac, BRD4, MED12, or RNAPII App. Part. of cells from (c). ***=p<0.0005, by two-sided unpaired T-test (n ≥ 428 cells, per condition). Line represents mean and error bars represent s.e.m. (e-f) Quantification of BRD4 or MED12 App. Part. of MCF7 (e) or HeLa (f) cells subjected to HS and siCtrl or siHSF1 siRNAs. ***=p<0.0005, by two-sided unpaired T-test (n ≥ 382 cells, per condition). Line represents mean and error bars represent s.e.m. (g) Representative pseudo-colored images of UHKO-HaH cells with or without dox. induction, subjected to HS, and stained for BRD4 or MED12 (red). Scale bar, 10 μm. Zoom-ins are 5 μm × 5 μm. (h) Quantification of BRD4 or MED12 App. Part. of cells from (g). ***=p<0.0005, by two-sided unpaired T-test (n ≥ 895 cells, per condition). Line represents mean and error bars represent s.e.m. (i) Representative pseudo-colored fluorescence images of U2OS cells treated with HS and DMSO, A-485, or OTX-015 and stained for HSF1 (green). Scale bar, 10 μm. Zoom-ins are 7.5 μm × 7.5 μm. (j) Quantification of HSF1 MFI, App. Part., mean foci area, and mean foci Ecc. in cells from (i). **=p<0.005, ***=p<0.0005, ns = not significant, by two-sided unpaired T-test (n ≥ 290 cells). Line represents mean and error bars represent s.e.m. (k) Meta-images of MED12, H3K27ac, BRD4, or RNAPII signal (red) across segmented HSF1 condensates (green) in cells subjected to HS and further treated with DMSO, A-485, or OTX-015. Intensity scale for respective biomolecules is also represented. (n ≥ 290 cells, 9599 condensates, per condition).

Considering that hubs are sensitive to epigenetic perturbation^37^, we tested if A-485^38^, a CBP/P300 inhibitor that impairs H3K27 acetylation (Supplementary Fig. 4a-b), or OTX-015^39^, an orally bioavailable BRD2/3/4 inhibitor, affected HS-induced HSF1 hubs. IF coupled 3D-HILO showed that HSF1 condensate formation persisted through these perturbations, exhibiting higher partition even in the absence of any significant change in HSF1 amount (Fig. 2i-j). Intriguingly, neither of these drugs affected MED12 co-enrichment at HSF1 condensates (Fig. 2k and Supplementary Fig. 4c), suggesting this aspect to be an early event in hub formation, that occurs independent of H3K27ac or BRDs. However, these HSF1 condensates had significantly lesser area and eccentricity as compared to DMSO controls (Fig. 2i-j), indicating that morphological changes may affect function. Corroborating this thesis, A-485 ablated the co-enrichment of H3K27ac, BRD4 and RNAPII, at HSF1 condensates. Given that BRD4 binds acetylated histones, reduction in H3K27ac concordantly diminishes BRD4 recruitment, and consequently BRD-interacting RNAPII recruitment, to HSF1 condensates. OTX-015 impaired co-condensation of BRD4 and RNAPII with minimal impact on H3K27ac levels or accumulation at HSF1 condensates (Fig. 2k and Supplementary Fig. 4d-f), consistent with impaired recruitment of BRD4-RNAPII complexes to chromatin. Overall, our data shows that HSF1 drives coordinated formation of HS-induced transcription hubs.

### Activation vs disorder in HSF1 hubs

HSF1 activation via extensive PTMs, trimerization and DNA binding are required for its function as a stress-responsive transcription factor during proteotoxic stress^12^. Thus, we tested if these factors impacted HSF1 condensate formation and function during HS. As expected for extensive PTMs, HSF1 migrated higher in immunoblots during HS as compared to unstressed conditions (Supplementary Fig. 5a). Phosphorylation of HSF1-S320 (pS320), a well validated proxy for HSF1 activation^40^, was significantly elevated during HS and (Supplementary Fig. 5a) and enriched within condensates (Fig. 3a). HSF1 inactivation by Cycloheximide(CHX)-induced translation inhibition^41^ significantly reduced HSF1 migration (almost to unstressed levels), pS320 levels (∼4-fold) and HSF1 condensate formation (∼4-fold, Fig. 3b and Supplementary Fig. 5b), strongly suggesting that activating PTMs drive HSF1 condensate formation. Using our prior established strategy of introducing Halo-HSF1 at physiological levels, without interference of endogenous HSF1 (UHKO-HaH, Fig. 2g-h and Supplementary Fig. 3l-n), we tested the role of distinct intramolecular regions in the form and function of HSF1 condensates via domain deletions (Fig. 3c). Using HTIHR, we found that removal of the DNA binding domain (UHKO-HaHΔDBD / ΔDBD) or the trimerization domain (UHKO-HaHΔLZ1-3 / ΔLZ1-3) ablated HSF1 condensate formation (Fig. 3d-e and Supplementary Fig. 5c-d) as compared to FL Halo-HSF1 (UHKO-HaH), even when expressed at similar levels (Supplementary Fig. 5e). Akin to the lack of HSF1 (Fig. 2c-f and Supplementary Fig. 3a-k), ΔDBD and ΔLZ1-3 lost the ability to function as BRD4-containing transcription hubs (Fig. 3f and Supplementary Fig. 5f). Concordantly, the topoisomerase poison Doxorubicin (Doxo) that evicts histones and TFs^42^ from DNA also ablated endogenous HSF1 condensate formation in U2OS cells (Fig. 3b). Removal of the HSF1 oligomerization repressor domain^12^ LZ4 (UHKO-HaHΔLZ4 / ΔLZ4), had minor effects on HSF1 condensation (partition, ∼1.25-fold reduction, Fig. 3d-e, and number per nucleus, 2-fold reduction, Supplementary Fig. 5c, as compared to FL), but no significant differences in the percentage of foci bearing cells (Supplementary Fig. 5d). Unlike ΔDBD and ΔLZ1-3, BRD4 can still be recruited to ΔLZ4 condensates, albeit to a lesser extent (∼2.5-fold as compared to FL, Fig. 3f and Supplementary Fig. 5f). Together, these data suggest that canonical activation is a requisite for HS-induced HSF1 condensate formation and function.

**Figure 3.**
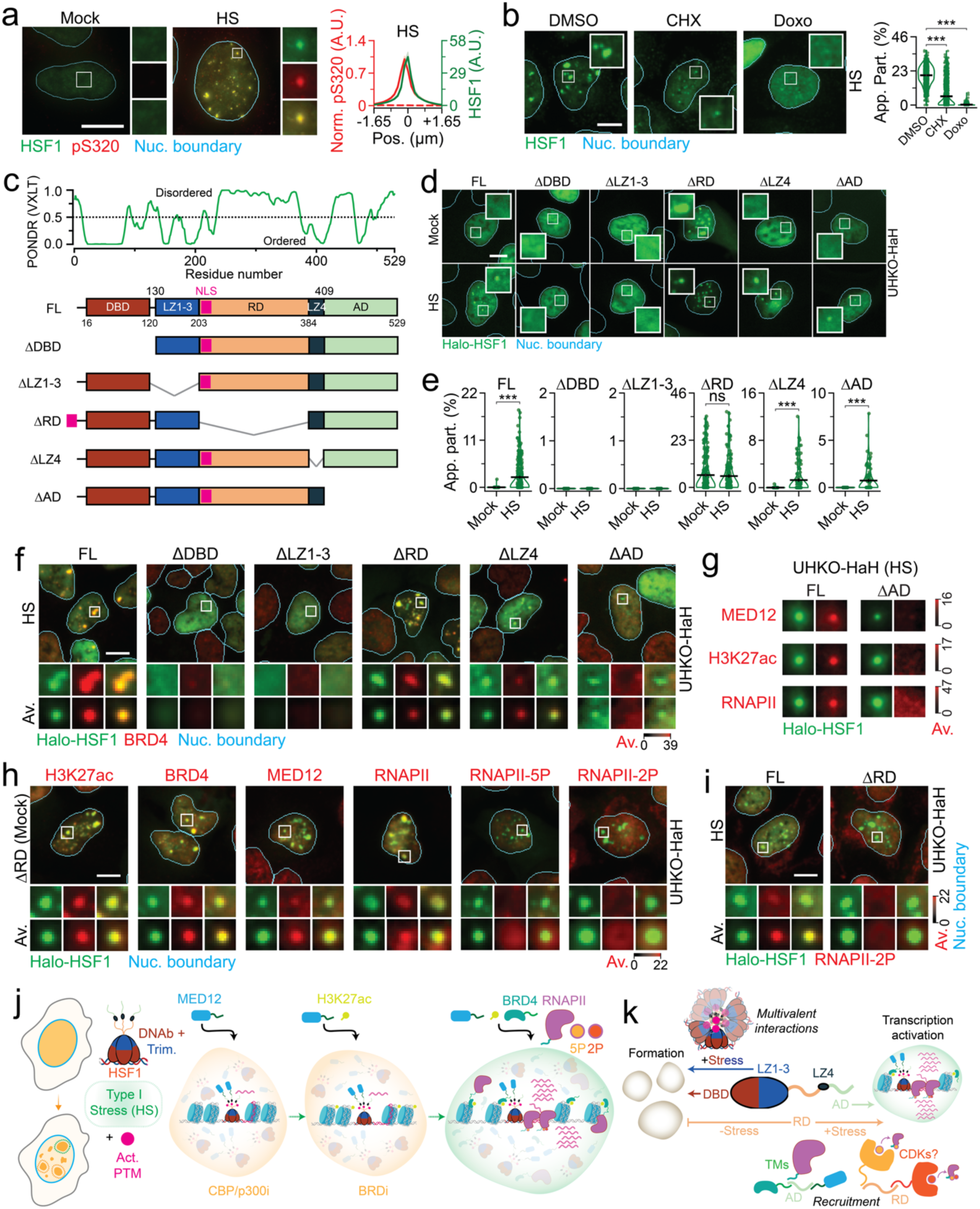
Activation and disorder modulate HSF1 condensate formation and function during HS. (a) Representative pseudo-colored IF image of U2OS cells treated with mock and HS conditions and stained for HSF1 (green) and HSF1-pS320 (pS320, red); with normalized line scan metaplot of HSF1-pS320 signal across segmented HSF1 condensates (n ≥ 94 cells, 2766 condensates, per condition). Dashed red lines represent co-IF signal from spatial randomizations. Scale bar, 10 μm. Zoom-ins are 2.7 μm × 2.7 μm. (b) Representative pseudo-colored IF images of HS subjected U2OS cells treated with DMSO, CHX, or Doxo; with quantification of HSF1 apparent partition (App. Part.). ***=p<0.0005, by two-sided unpaired T-test (n ≥ 900 cells per condition). Line represents mean and error bars represent s.e.m. Scale bar, 10 μm. Zoom-ins are 12.5 μm × 12.5 μm. (c) Disorder prediction from PONDR along with schematic of HSF1 domain deletion constructs. (d) Representative pseudo-colored fluorescent images of mock or HS treated FL, ΔDBD, ΔLZ1-3, ΔRD, ΔLZ4, and ΔAD cells stained with Halo-HSF1 (green). Scale bar, 10 μm. Zoom-ins are 7.5 μm × 7.5 μm. (e) Quantification of Halo-HSF1 App. Part. in from cells in (d). ***=p<0.0005, by two-sided unpaired T-test (n ≥ 300 cells, per condition). Line represents mean and error bars represent s.e.m. (f) Representative pseudo-colored images of HS subjected FL, ΔDBD, ΔLZ1-3, ΔRD, ΔLZ4, and ΔAD cells stained with Halo-HSF1 (JF549-HL, green) and BRD4 (red); with meta-images (Av., n ≥ 69 cells, per condition). Scale bar, 10 μm. Zoom-ins and meta-images are 4.2 μm × 4.2 μm. (g) Meta-images of MED12, H3K27ac, or RNAPII signal (red) across segmented Halo-HSF1 condensates (JF549-HL, green) in FL, or ΔAD cells subjected to HS. Intensity scale for respective biomolecules is also represented. (n ≥ 67 cells, 436 condensates, per condition). (h) Representative pseudo-colored images of mock-treated ΔRD cells stained with Halo-HSF1 (JF549-HL, green) and H3K27ac, BRD4, MED12, RNAPII, RNAPII-5P, or RNAPII-2P (red); with meta-images (Av., n ≥ 644 cells, per condition). Scale bar, 10 μm. Zoom-ins and meta-images are 4.2 μm × 4.2 μm. (i) Representative pseudo-colored images of heat-shocked FL and ΔRD cells stained with Halo-HSF1 (JF549-HL, green) and RNAPII-2P (red); with meta-images (Av., n ≥ 284 cells, per condition). Scale bar, 10 μm. Zoom-ins and meta-images are 4.2 μm × 4.2 μm. (j) Model representing coordinated hub formation during HS. (k) Model representing *cis* elements and *trans* factors that drive HSF1 condensate formation and function during HS.

IDRs are foundational drivers of condensate formation^10^. However, removal of HSF1’s disordered regulatory domain (UHKO-HHΔRD / ΔRD) spontaneously induced condensate formation in the absence of stress (Fig. 3c-e and Supplementary Fig. 5c-d). Moreover, HSF1 lacking the significantly disordered activating domain (UHKO-HHΔAD / ΔAD), which engages chromatin remodeling factors and RNAPII^43^, still reorganized from a dispersed (unstressed) to a condensed (during HS) state, albeit to a lesser extent (∼2.5-fold reduction than FL, Fig. 3c-e and Supplementary Fig. 5c-d). However, ΔAD condensates were unable to coenrich for BRD4, MED12, H3K27ac and RNAPII (Fig. 3f-g and Supplementary Fig. 5f-h), underscoring the need for this region for transcriptional competency. On the other hand, ΔRD condensates co-enriched for BRD4 in the presence (Fig. 3f and Supplementary Fig. 5f) and absence (Fig. 3g and Supplementary Fig. 5i) of HS and enrich for almost all the transcription machinery (H3K27ac, MED12 and RNAPII) under unstressed conditions (Fig. 3h and Supplementary Fig. 5i). But ΔRD condensates were devoid of RNAPII-5P and RNAPII-2P in unstressed conditions (Fig. 3h and Supplementary Fig. 5i) and these condensates still lacked RNAPII-2P during HS (Fig. 3h and Supplementary Fig. 5j). In essence, disordered regions are not needed for HSF1 condensate formation *in situ*, with the RD regulating condensate formation. However, HSF1’s IDRs are necessary for condensate maturation to transcription hubs during HS.

Using chemical and genetic perturbations (Fig. 2-3) we have generated snapshots of HS-induced hubs (Type I) at distinct stages of activation and unraveled factors required for condensate formation and function (Fig. 3j-k and Supplementary Fig. 5k-l). Our data suggests an orchestrated hub assembly in human cells (Fig. 3j), wherein activated HSF1 recruits MED12, H3K27ac is potentially remodeled at these co-condensates, acetyl-binding BRD4 is then co-enriched, which in turn further recruits RNAPII. Here, structured HSF1 *cis* elements (DNA binding and trimerization domains) and PTMs drive HSF1 condensate formation via initial multivalent interactions^1^, while disordered HSF1 regions serve to recruit and enrich *trans* factors for transcriptional function (Fig. 3k). Specifically, the disordered AD, in line with its transactivation function across TFs, recruits transcriptional cofactors via its proposed co-condensation capacity^44^ (Fig. 3k and Supplementary Fig. 5k-l). The disordered RD prevents condensate formation in the absence of stress, but is required to transition cofactor-enriched condensates to transcription hubs, possibly via recruitment of factors that mediate CTD phosphorylation^32^ (Fig. 3k and Supplementary Fig. 5k-l).

### Aberrant hub organization in stresses

Having established the mechanisms of HSF1 condensate formation and function during HS, we sought to understand the molecular logic behind diminished transcriptional competence of HSF1 condensates in other stresses. Meta-analysis of nano-to-mesoscale signal distribution across thousands of HSF1 condensates showed that the full complement of factors required for functional hub formation was lacking in all other stresses (Fig. 4a and Supplementary Fig. 6a-j). Specifically, all components of the transcription apparatus including H3K27ac, BRD4, MED12, RNAPII and RNAPII-5P were enriched at HSF1 condensates that form during ACA toxin induced proteotoxic stress, yet these condensates had little-to-no RNAPII-2P. HSF1 condensates that form during CdC-induced heavy metal toxicity contained H3K27ac, BRD4 and MED12, but not RNAPII, and consequently no RNAPII-5P and RNAPII-2P. Oxidative stress (SA and HP) driven HSF1 condensates largely contained MED12 only. HSF1 condensates formed during CC-induced combined chemical hypoxia and heavy metal stress recruited BRD4 and MED12 only. During MGO-induced metabolic stress, HSF1 condensates contained BRD4 only, with chemotherapeutic (ATO) induced HSF1 condensates not enriching for any of the transcription machinery.

**Figure 4.**
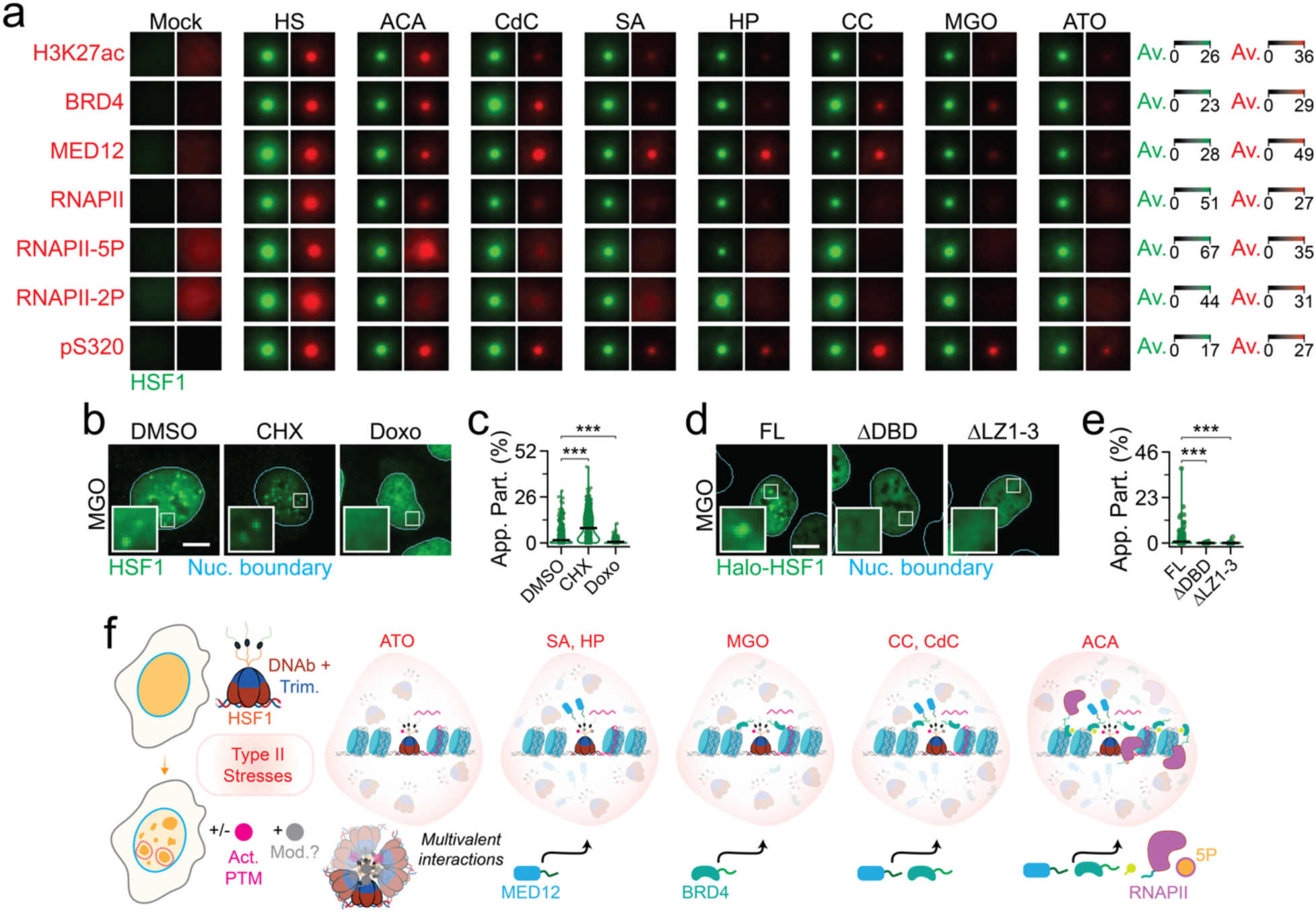
PTM-agnostic HSF1 condensates are stalled at distinct stages of hub formation during other stresses. (a) Meta-images of H327ac, BRD4, MED12, RNAPII, RNAPII-5P, RNAPII-2P, and HSF1-pS320 (pS320) in U2OS cells treated with mock, HS, ACA, CdC, SA, HP, CC, MGP, or ATO conditions across segmented HSF1 condensates (n ≥ 53 cells, 1094 condensates, per condition). Meta-images are 2.7 μm × 2.7 μm. (b) Representative pseudo-colored IF images of U2OS cells treated with MGO, along with DMSO, CHX, or Doxo, and stained for HSF1 (green). Nuc. Boundary (cyan) is derived from DAPI staining. Scale bar, 10 μm. Zoom ins are 5 μm × 5 μm. (c) Quantification of HSF1 App. Part. in from cells in (b). ***=p<0.0005, by two-sided unpaired T-test (n ≥ 409 cells, per condition). Line represents mean and error bars represent s.e.m. (d) Representative pseudo-colored fluorescent images of FL, ΔDBD, and ΔLZ1-3 cells treated with MGO and stained for Halo-HSF1 (JF549-HLgreen). Nuc. Boundary (cyan) is derived from DAPI staining. Scale bar, 10 μm. Zoom ins are 5 μm × 5 μm. (e) Quantification of HSF1 App. Part. in from cells in (d). ***=p<0.0005, by two-sided unpaired T-test (n ≥ 120 cells, per condition). Line represents mean and error bars represent s.e.m. (f) Model of transcriptionally incompetent HSF1 condensates, lacking a full complement of factors required for functional hubs, in other stresses.

Since activation is a pre-requisite for HS-induced transcription hubs, we then checked if this aspect was impacted in these stresses. Indicative of distinct types of PTMs acquired by HSF1, its migration pattern was significantly different between HS and all other stresses (Supplementary Fig. 7a). While pS320 levels in most other stresses, except of MGO and ACA, were significantly higher than mock, they were much lower when compared to HS (Supplementary Fig. 7a), suggesting insufficient activation. Concordantly, pS320 enrichment at HSF1 condensates of other stresses were significantly lower than that of HS (reduction by ∼1.5-fold in ACA, ∼1.8-fold in CdC, ≥ 10-fold in CdC, SA, HP, MGO and ATO, and ∼1.3-fold in CC, as compared to HS, Fig. 4a and Supplementary Fig. 7b-c). However, treatment with HSF1 PTM-inhibiting CHX enhanced endogenous HSF1 condensate formation (∼1.3 – 5-fold) during ATO, CdC, SA and MGO stresses (Fig. 4b-c and Supplementary Fig. 7d-e) as compared to CHX dissolving HS-induced HSF1 condensates (Fig. 3b). Moreover, ΔDBD, DNA binding mitigating Doxo treatment, and ΔLZ1-3 exhibited dispersed signal distribution during non-HS stresses (Fig. 4b-e, and Supplementary Fig. 7d-g). Together, our data suggest that HSF1 condensates form independent of activating PTMs during many other (Type II) stresses, but these condensates are still dependent on DNA binding and trimerization of HSF1 (Fig. 4f). Depending on the stress-type, HSF1 condensates are stalled at distinct stages of hub formation (Fig. 4f), underscoring diverse modes of condensate dysfunction.

### Condensate defects curb HSF1 loci output

Biomolecules within functional and aberrant condensates have been reported to exhibit distinct physicochemical properties^8^. To test this concept, we developed an assay for intra-condensate single-molecule tracking (SMT, Fig. 5a and Supplementary Video 1). Analysis of individual tracks and diffusion constants showed that HSF1 molecules were relatively more dynamic (∼5 – 10-fold) within HS-induced HSF1 condensates, as compared to condensates in other stresses (Fig. 5b-d). Thus, type I and type II HSF1 condensates potentially have distinct material properties. We then employed single-molecule RNA fluorescence *in situ* hybridization (smFISH) to quantitatively assess the impact of type I and II stresses on transcriptional output HSF1-regualted genes both within and outside of HSF1 condensates. IF-smFISH showed that HSATIII non-coding RNAs were induced in HS (9.5-fold), SA (2.4-fold) and HP (2.1-fold) stresses (Fig. 5e-f). Strikingly, the extent of HSATIII enrichment at SA- and HP-induced HSF1 condensates were 10 – 100-fold lesser than HS, with HSATIII induction not occurring within HP-induced condensates (Fig. 5g). SmFISH analysis of HSP90AA1, a chaperone locus typically found within HSF1 nanoclusters^21^, showed that the gene is expectedly induced at its transcription site (Fig. 5h) during HS and SA, but the extent of induction is ∼5-fold higher in HS, with no induction during HP stress (Fig. 5i-j). These data indicate a correlation between dynamic condensates and transcription at these locations and suggest that HSF1-mediated muted gene induction occurs outside of nano-to-mesoscale HSF1 condensates.

**Figure 5.**
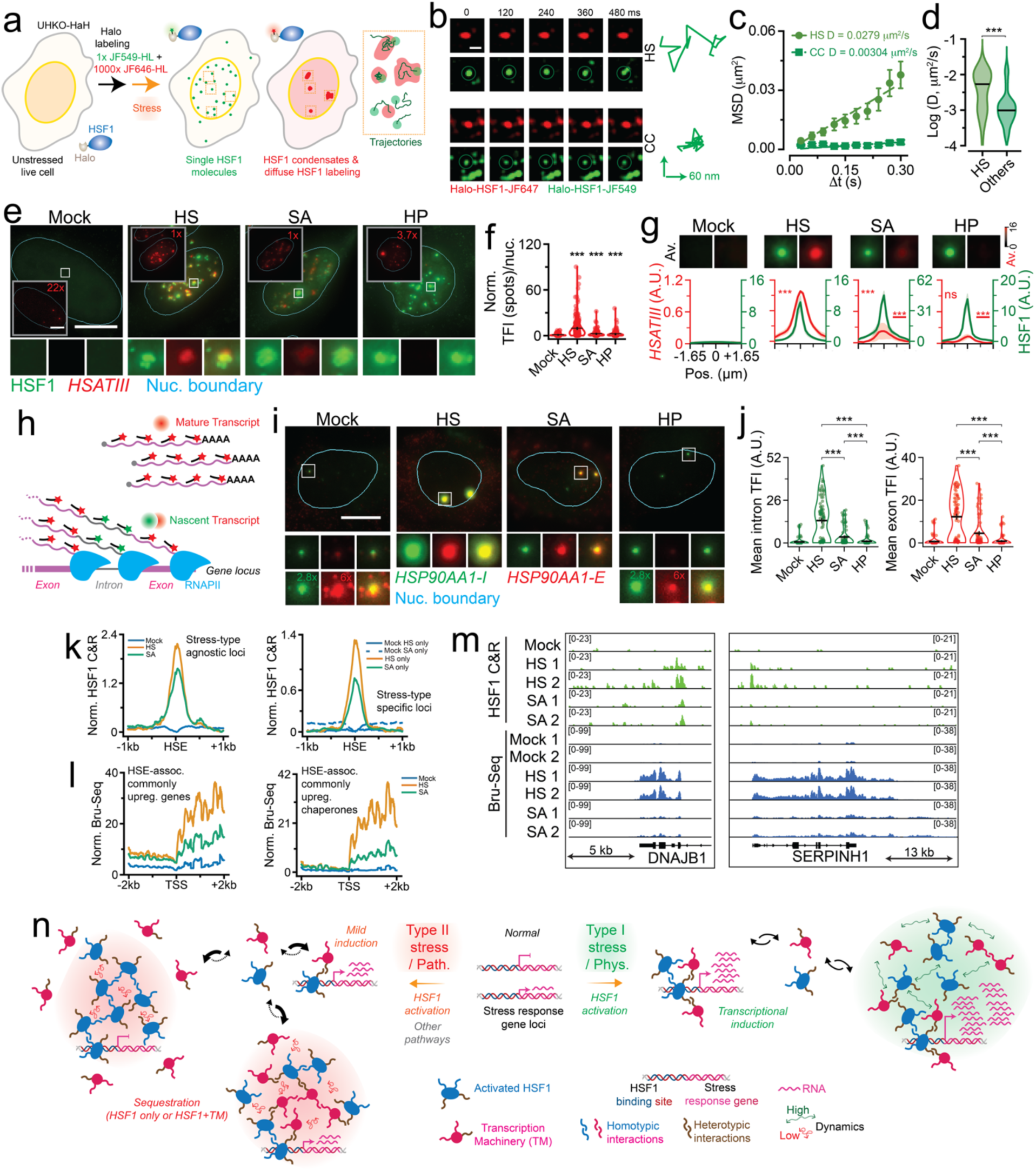
Sequestration of HSF1 is associated with impairment of HSF1-mediated transcriptional stress response. (a) Schematic of intra-condensate SMT assay. (b) Representative pseudo-colored time-lapsed images of HSF1 condensates (JF646-HL, red) and single-HSF1 molecules (JF549-HL, green) from live UHKO-HaH cells subjected to HS or CC. Dotted green circles have been included to aid HSF1 molecule location. Appropriate single-HSF1 molecule trajectory are also represented. Scale bar, 60 nm. (c) Mean square displacement (MSD) vs time-lag (τιt) plots for trajectories in (b). Diffusion coefficients (D) extracted from plots are indicated. (d) Distribution of log D of HSF1 molecules in HS and other stresses (Others: CC, HP, MGO, CdC, and ACA). ***=p<0.0005, by two-sided unpaired T-test (n ≥ 20 cells, 102 trajectories, per condition). Line represents median. (e) Representative pseudo-colored images of U2OS cells mock treated or subjected to HS, SA, or HP, and stained for HSF1 (green, by IF) and *HSATIII* (red, by smFISH); with meta-images (Av., n ≥ 58 cells, per condition). Nuc. Boundary (cyan) is derived from DAPI staining. Scale bar, 10 μm. Meta-images and zoom-ins are 2.7 μm × 2.7 μm. Inset represents smFISH signal, with distinct (x-fold) contrast adjustment to aid visualization. (f) Quantification of *HSATIII* signal at in cells from (e). ***=p<0.0005, by two-sided unpaired T-test. Line represents mean and error bars represent s.e.m. (g) Meta-images (Av.) and normalized line scan metaplots of *HSATIII* smFISH signal across segmented HSF1 condensates from (e). ***=p<0.0005, ns = not significant, by two-sided unpaired T-test (n ≥ 58 cells, 1750 condensates, per condition). Underscored significance denotations are relative to HS, whereas others are relative to Mock. (h) Schematic of transcription site (TS) imaging by smFISH. (i) Representative pseudo-colored smFISH images of U2OS cells mock treated or subjected to HS, SA, or HP, and stained for HSP90AA1 introns (green) or HSP90AA1 exons (red). Nuc. Boundary (cyan) is derived from DAPI staining. Scale bar, 10 μm. Zoom-ins are 2.7 μm × 2.7 μm. (j) Quantification of *HSP90AA1* intron and exon signal at putative TS in cells from (h). ***=p<0.0005, by two-sided unpaired T-test (n ≥ 80 cells, per condition). Line represents mean and error bars represent s.e.m. (k) Metaplot of σ; 120 bp fragments HSF1 CUT&RUN signal at HSEs in U2OS cells that were mock treated or subjected to HS, or SA. HSEs at stress-type agnostic (left) or stress-type specific (right) loci with upregulated signal are represented. (l) Metaplot of BrU-Seq signal in U2OS cells that were mock treated or subjected to HS, or SA. HSE associated commonly (stress-type-agnostic) upregulated genes (left) or chaperones (right) are represented. (m) Representative genomic view of DNAJB1 and SERPINH1 loci with HSF1 CUT&RUN (C&R) or BrU-seq signal from U2OS cells that were mock treated or subjected to HS, or SA. (n) Proposed model of divergent functionality of HSF1 and potentially other stress-induced transcription factor condensates.

Our data suggest that HSF1 condensates aberrantly form and function at DNA during a variety of non-HS stresses, including SA treatment (Fig. 4). To test address this aspect genome-wide during HS and SA stress, we performed fixation-free epigenomics via cleavage under targets and release under nuclease (CUT&RUN, Methods) to assess HSF1’s genomic occupancy and nascent RNA sequencing via metabolic RNA labeling with bromouridine (BrU-Seq, Methods) to assess transcriptional outcome. As previously reported for these stresses in other cell lines^14^, HSF1 localized to sites that were common to both stresses and to stress-type-specific sites in U2OS cells (Supplementary Fig. 8a). Leveraging non-human spike-ins to account for global changes in genomic occupancy, we found that the amount of HSF1-associated genomic reads was ∼2-fold higher in HS as compared to both mock and SA treatment (Supplementary Fig. 8b). Consistently, spike-in corrected HSF1 occupancy showed that HSF1’s binding at stress-type-agnostic and - specific HSEs were higher during stress when compared to unstressed controls, but net binding in HS was significantly higher than SA (Fig. 5k). BrU-Seq complemented CUT&RUN and showed that HS and SA treatment differentially regulated both stress-type-agnostic and stress-type-specific transcripts (Supplementary Fig. Fig 8c-d). Meta-analysis of HSF1-proximal commonly upregulated genes (Fig. 5l, left, and Supplementary Fig. Fig 8c-d) or chaperones (Fig. 5l, right), confirmed that transcriptional output of putative HSF1-regulated genes is higher in HS as compared to SA, as correlated with differences in HSF1 cistromic occupancy (Fig. 5m). Since many stress-type-agnostic transcripts exhibited higher or similar induction in SA when compared to HS (Supplementary Fig. Fig 8e-f), gene induction ability is still robust in SA, indicating that the differences we observe are not due to generic deficiencies in transcription. Together with SMT and smFISH, our data supports the notion that sequestration of HSF1 and transcription machinery within aberrant, DNA-bound, non-dynamic HSF1 condensates affects global genomic occupancy and transcriptional output in SA and potentially in other non-HS stresses.

## DISCUSSION

In essence, we delineate the *cis* elements and *trans* factors that mediate formation and function of HSF1 condensates *in situ* and unravel the genome-wide regulatory implications of condensate functionality across a variety of stressful scenarios. In physiological scenarios (Type I), homotypic multivalent interactions^1^ via structured *cis* elements and activating PTMs (e.g. phosphorylation, contributing via electrostatics) initiate condensation, with heterotypic interactions between HSF1 and *trans* factors (the transcription machinery, TM) potentially enhance condensation, driving transcriptional activity via disordered regions (Fig. 5n). Here, the concept of orchestrated hub assembly is akin to step-wise events in yeast^45^, but the progression is vastly different in humans. Intriguingly, we find that the disordered human RD is a negative regulator of condensate formation in the absence of stress. With the RD containing a majority of putative (PhosphoSitePlus®) and validated PTM sites^21^, changes in the PTM landscape of this domain likely serves as a switch that facilitates multivalency, possibly driving intramolecular rearrangement of structured domains that enhances intermolecular interactions. Together, our data aligns with observations of HSF1 accumulating within HS-induced transcriptional condensates in yeast^28,45^. Thus, HSF1 condensate function is evolutionary conserved during physiological stresses, such as hyperthermia, febrility and inflammation^18,46^, wherein hubs dynamically and robustly induce transcriptional responses (Fig. 5n).

HSF1 has been reported to bind promoters, enhancers^14,47^ and repetitive elements^48^. As DNA binding drives condensate formation, our nano-to-mesoscale Type I foci are likely to be at these genomic regions. However, the resolution of HILO (∼50 nm) is limiting and our condensates lack H3K4me3 enrichment, therefore we cannot visualize HSF1 nanoclusters (∼10-20 nm) at H3K4me3-marked promoters of chaperone genes^21^. A significant minority of our condensates potentially represent mesoscale (primary and secondary) nSBs that associate with pericentromeric regions^48^. Considering that only a small fraction (∼17%) of our condensates enrich for HSATIII and our nano-to-mesoscale HSF1 foci copy number (∼15-50) far exceeds that of nSBs (∼1-12) from prior reports^48^, it is possible that a majority of our nanoscale HSF1 condensates are at other genomic locations, such as enhancers. Regardless of genomic location, Type I condensates represent a functional class of loci with enhanced transcription. In yeast, HS-induced HSF1 condensates have been reported to reorganize the genome^28^, wherein genes located on different chromosomes coalesce for enhanced transcription, but such large scale chromatin conformation changes seem to be absent in heat-shocked human cells^47^. Considering that many HSF1 dependent genes exist within the same topologically associated domain in human cells^47^, a single HSF1 condensate may induce multiple genes, akin to the functional outcome in yeast. Primate-specific mechanisms could include HSF1-bearing nSBs inducing the expression of 3D proximal genes^18^. Finally, factors that have been shown to drive HSF1’s TF activity^12^ are required for hub formation and HSF1 nanoclusters^21^. These observations suggest that nuclear HSF1 is spread across a continuum of dispersed-to-condensed states during HS, with the extent of condensation correlating with that of transcription induction to mete out loci-specific output during HS (Fig. 5n).

During Type II stresses (ATO, MGO, SA, HP, CC, CdC and ACA), HSF1 condensation still requires DNA binding and trimerization, underscoring their contribution to multivalency. However, HSF1 condenses form independent of activating PTMs, suggesting that canonical activation is triggered but additional mechanisms drive HSF1 sequestration within transcriptionally impaired condensates. These alternate mechanisms potentially include glycation (by MGO); toxic metalloids and metals (by As from SA and ATO, and Cd from CdC) chelating specific amino acids, especially thiol coordination; oxidative damage of side chains (by SA, HP and CC) and incorporation of unnatural amino acids (proline-like ACA). All these stresses are generally proteotoxic^14–16,19,31^ and consistently result in chaperone gene induction^14–16,19^. Our data showing the presence of activating PTMs on HSF1, albeit to minimal levels, aligns with these findings. Since HSF1 contains many of these modifiable amino acids (e,g, Arginine, Lysine, Cysteine and Proline), all these stresses likely impact the protein itself and promote its sequestration into dysfunctional condensates. Notably, the extent of condensation within transcriptionally inactive condensates correlates with extent of reduction in cistromic occupancy and transcriptional output (Fig. 5e-n). Our findings rationalize therapeutic and pathological scenarios, wherein the central arbiter of proteostasis is disrupted. For instance, ATO’s anticancer effects are potentially enhanced by sequestering HSF1 within HSR-incompetent aggregates and consequently inducing cytotoxicity - a new avenue for developing anticancer therapies. On the other hand, build-up of toxic metabolites like MGO (frequently occurs in diabetes and aging-related diseases), increased reactive oxygen species (as observed in neurodegenerative diseases and male infertility), and acute exposure to environmental pollutants, sequester HSF1 or distinct stress-type specific transcription apparatus into HSF1-depednent inclusions, essentially dampening the cytoprotective HSR (Fig. 5n). In this manner, our work highlights expression-independent HSF1 dysfunction as a contributor to the pathology of degenerative diseases and acute toxicities, in addition to prior reports of pathology-associated changes in HSF1 levels^23^. Additionally, transcriptional apparatus sequestration mechanistically rationalizes anti-correlation between large HSF1 foci and chaperone levels in tumor resections^27^, suggesting that diffuse-to-nanoscale condensates may mediate non-oncogene addiction to HSF1^24^. Overall, our study underscores the note that TF condensate formation does not predict increased transcriptional activity and contextualizes the contribution of HSF1 condensates in physiology and pathology.

## METHODS

### Plasmid construction

pLVX-TetOne-Puro-Halo-GGSx3-MCS was created from modifications of pLVX-TetOne-Puro-hAXL (Addgene #124797). The endogenous hAXL coding sequence was removed and replaced with a synthesized gene block (Twist Bioscience®) encoding a HaloTag, GGSx3 linker sequence and a multiple cloning site (MCS). pLVX-TetOne-Puro-Halo-GGSx3-HSF1 was created by PCR amplification of the HSF1 coding sequence from HSF1-GFPN3 (Addgene #32538) and subsequently cloned into the MCS using AgeI and NotI restriction sites. pLVX-TetOne-Puro-Halo-GGSx3-HSF1-ΔDBD, pLVX-TetOne-Puro-Halo-GGSx3-HSF1-ΔLZ1-3, pLVX-TetOne-Puro-Halo-GGSx3-NLSx3-HSF1-ΔRD, pLVX-TetOne-Puro-Halo-GGSx3-HSF1-ΔLZ4 and pLVX-TetOne-Puro-Halo-GGSx3-HSF1-ΔAD were created by cloning in synthesized gene blocks (Twist Bioscience®) of HSF1-ΔDBD, HSF1-ΔLZ1-3, NLS3x-HSF1-ΔRD, HSF1-ΔLZ4, HSF1-ΔAD into pLVX-TetOne-Puro-Halo-GGSx3-MCS. Since HSF1’s NLS is within the RD, we added 3x SV40-NLS to the N-terminal of HSF1-ΔRD construct. All sequences were verified by whole plasmid sequencing (Plasmidosaurus®).

### Cell culture and stressor/drug treatment

Human osteosarcoma U2OS cells (ATCC, HTB-96) were maintained in McCoy’s 5A modified medium (Gibco, 16600108) contains 10% fetal bovine serum (FBS, Gibco, A5256501) and 100 U/mL Penicillin-Streptomycin (P-S, Gibco, 15140122). Human cervical carcinoma, HeLa (ATCC, CCL-2) and human breast carcinoma MCF7 (ATCC, HTB-22) were maintained in DMEM (Gibco, 11995-065) supplemented with GlutaMAX (Gibco, 3050061), 10% FBS and 100U/mL P-S. Cells were cultured in at 37 °C and 5% CO_2_. For heat shock (HS) cells were transferred to a 42 °C incubator with 5% CO_2_ for 1 h. All other stress treatments were performed by supplementing the appropriate stressor into cell medium and incubation at 37 °C with 5% CO_2_. Stressor treatments were as follows: Sodium arsenate (SA, Sigma Aldrich, S7400, 0.5 mM, 1 h; Hydrogen peroxide (HP, Sigma Aldrich, 216763, 2 mM, 1 h; Arsenic trioxide (ATO, Sigma Aldrich, A1010, 0.5 mM, 1 h; Cobalt chloride (CC, Sigma Aldrich, C8661, 10 mM, 4 h; Cadmium chloride (CdC; Sigma Aldrich, 202908 1 mM, 1 h; Methylglyoxal (MGO, Sigma Aldrich, M0252) 10mM, 1 h; L-azatidine-2-carboxylic acid (ACA, Sigma Aldrich, A0760 10 mM, 6 h. All drug treatments were performed by pre-incubating cells with the appropriate drugs for 4 h at 37 °C with 5% CO_2_ and then co-treatment with the stressor. Drug treatments were as follows: A-485 (MedChemExpress, HY-107455), 100 µM; OTX-015 (MedChemExpress, HY-15743), 100 µM; Cycloheximide (CHX, Sigma Aldrich, C7698),10 µg/mL; and Doxorubicin (Doxo, MedChemExpress, HY-15142), 10 µM.

### Generation of HSF1 knockout cells

U2OS cells lacking HSF1 (UHKO) were created using CRISPR/Cas9 mediated gene editing. Briefly, guide RNAs (sgRNAs) targeting the exons of human *HSF1* were designed using Benchling. Non-targeting sgRNA and *HSF1*-targeting sgRNAs were cloned into lentiCRISPR v2 plasmid (Addgene plasmid #52961). U2OS cells were transiently transfected with lentiCRISPR v2 encoding non-targeting or pool of three independent HSF-targeting sgRNAs. Two days after transfection, cells were selected with 1 μg/ml puromycin (ThermoFisher, A1113803) for three days. Western blotting (below) was performed to examine the knock-out efficiency, then single cells were seeded into 96-well plates to get monoclonal UHKO cells, which were cultured akin to the parental U2OS cells. The sgRNA sequences are listed in Supplementary Table 1.

### HSF1 knockdown

*HSF1* gene silencing was achieved using ON-TARGETplus Human HSF1 siRNA SMARTpool (siHSF1, Horizon Discovery, L-012109-02-0005). Control siRNAs (siCtrl, Thermo, 4457287) were used for comparisons. siRNAs were transfected using Lipofectamine RNAiMAX (Invitrogen, 13778150) in 6-well polystyrene plates. HSF1 knockdown was checked 48 h after transfection by western blotting. For imaging assays that required HSF1 knockdown, siCtrl or siHSF1 transfected cells were trypsinized from 6-well plates 24 h after transfection and reseeded into either 8 well chambered cover glass (Cellvis, C8-1.5H-N) or 96 well glass bottom plate (Cellvis, P96-1.5H-N). After an additional 24 h incubation, cells were processed for imaging.

### Generation of inducible knock-in cells and doxycycline (dox.) induction

Dox. inducible knock-in of full-length (FL) Halo-HSF1 (HaH) or different domain deletions were created by stably transducing UHKO cells with various constructs embedded in the pLVX-TetOne-Puro-Halo-GGSx3-MCS backbone. Briefly, UHKO cells were transduced with the appropriate lentivirus for 12 h. The viral supernatant was then removed and replaced with fresh medium for 48 h. Cells were subsequently subjected to antibiotic selection using puromycin (1 µg/mL), encoded by the lentiviral vector. Selection medium was replaced every 2 days and continued for 7 days until all non-transduced control cells were eliminated. Puromycin-resistant cells were expanded and maintained in U2OS growth medium supplemented with a low dose of puromycin (250 ng/mL) for downstream experiments. The following cells were created, UHKO-HaH (FL), UHKO-HaH-ΔDBD, UHKO-HaH-ΔLZ1-3, UHKO-HaH-ΔRD, UHKO-HaH-ΔLZ4, UHKO-HaH-ΔAD. To induce expression of HaH (FL) or HaH domain deletions, appropriate knock-in cells were treated with 0.1 µg/mL dox. (Sigma, D3447) for 24 h. Expression after dox. induction was checked by western blotting.

### Fluorescent labeling and processing of knock-in cells for condensate imaging

For fixed cell imaging of dox. induced knock-ins, cells that were seeded and induced in 8 well chambered cover glass or 96 well glass bottom plate were labeled with 200 nM JF549-HL (gift from the Lavis Lab, JF549 Halotag ligand) diluted in growth medium for 1 h. Followed by 2x 15 min washes with growth medium, cells were washed 2x with pre-warmed (37°C typically or 42°C for HS) 1x PBS (Gibco, 10010031) and immediately fixed with prewarmed (37°C or 42°C as appropriate) fixation buffer (4% paraformaldehyde/PFA in 1x PBS; PFA, Electron Microscope Sciences, SKU15710) for 10 min at room temperature (RT). Cells were then washed 3x with 1x PBS and permeabilized with permeabilization buffer (0.5 % Triton-X-100 in 1x PBS; Triton-X-100, Sigma, T8787) for 10 mins at RT and washed 2x with 1x PBS. Cells were counterstained with 50 ng/mL DAPI (Sigma, 9542) with or without 0.5 µg/mL NIR-Cell Mask (Invitrogen, H32722) in 1x PBS for 10 min at RT in the dark. Finally, cells were washed 2x with 1x PBS. Cells were mounted in PBS antifade (PBS-TPP, 1mM Trolox, 4.5 mM 3,4-Dihydroxybenzoic acid/PCA and 0.135 U/mL protocatechuate dehydrogenase/PCD in 1x PBS; Trolox, Acros Organics, 8823395; 3,4-Dihydroxybenzoic acid, Sigma, 37580; protocatechuate dehydrogenase, MedChemExpress, HY-P2915) and overlaid with low density mineral oil (Sigma, 330779) prior to imaging.

### Immunofluorescence (IF)

Cells in 8 well chambered cover glass or 96 well glass bottom plate were washed 2x with pre-warmed (37°C or 42°C) 1x PBS and immediately fixed with prewarmed (37°C / 42°C) fixation buffer (4% PFA in 1x PBS) for 10 min at RT. Cells were then washed 3x with 1x PBS and permeabilized with permeabilization buffer (0.5 % Triton-X-100 in 1x PBS) for 10 mins and washed 2x with 1x PBS. Blocking was performed using blocking buffer (BB, 2.5% normal goat serum in 1x PBS; normal goat serum, Jackson Immuno Research, 005-000-121,) for 1 h at RT. Cells were incubated with primary antibodies (pAs) in BB for 1 h, followed by 3x 5 min washes with BB. Cells were then incubated with fluorophore conjugated secondary antibodies (sA) in BB for 1 h at RT in the dark. Subsequently, cells were subjected to 2x 5 min washes with BB and 1x 5 min wash with 1x PBS. Cells were counterstained with 50 ng/mL DAPI with or without 0.5 µg/mL NIR-Cell Mask in 1x PBS for 10 min at RT in the dark. Finally, cells were washed 2x with 1x PBS. Cells were mounted in PBS antifade (PBS-TPP, 1mM Trolox, 4.5 mM 3,4-Dihydroxybenzoic acid and 0.135 U/mL protocatechuate dehydrogenase in 1x PBS) and overlaid with low density mineral oil prior to imaging. pAs used were as follows: rb-a-HSF1 (abcam, ab52757), rat-a-HSF1 (abcam, ab61382), mmu-a-HSF1 (Santa Cruz Biotechnology, sc-17757), rb-a-H3K4Me3 (Cell Signaling Technology, C42D8), rb-a-H3K27Ac (Sigma, HPA061646), rb-a-MED12 (Cell Signaling Technology, 14360S), mmu-a-RNAPII (Diagenode, C15200004), mmu-a-RNAPII-2P (Diagenode, C15200005), mmu-a-RNAPII-5P (Diagenode, C15200007), rb-a-HSF1-phospo S320 (abcam, ab76183). Secondary antibodies (sAs) used were as follows: goat-a-rb-Cy3 (Jackson Immuno Research, 111-165-144), goat-a-rb-Cy5 (Jackson Immuno Research, 111-175-144), goat-a-rat-cy3 (Jackson Immuno Research, 112-165-167), goat-a-mmu-cy5 (Jackson Immuno Research, 115-175-165). For IF assays that used dox. induced, JF549-HL labeled, knock-ins, cells were processed for IF right after the 2x 15 min washes with growth medium.

### Western blotting (WB)

Cells on 6-well plates were washed 2x with ice-cold 1x PBS and lysed on ice in RIPA buffer (Thermo Fisher Scientific, 89901; ∼200 µL per well) supplemented with 1× protease and phosphatase inhibitors (Halt Protease Inhibitor Cocktail, 100×, 78430). Lysates were scraped, collected into centrifuge tubes, sonicated (6 s pulse, 20 s pause, 35% amplitude), and centrifuged at 12,000 rpm for 10 min at 4 °C. Supernatants were collected and stored at −80 °C. Protein concentrations were determined using the Pierce™ BCA Protein Assay with BSA standards. Protein (10 µg) were mixed with NuPAGE LDS Sample Buffer (4×, NP0007) and NuPAGE Sample Reducing Agent (10×, NP0004) and denatured at 95 °C for 10 min. Samples were resolved on NuPAGE 4–12% Bis-Tris gels using MES running buffer (100 V for 15 min, followed by 125 V for 45 min). A Spectra Multicolor Broad Range Protein Ladder (Thermo Fisher Scientific; 26634) was run in parallel. Proteins were transferred onto PVDF membranes using a semi-dry transfer system (Bio-Rad Trans-Blot Turbo Mini 0.2 µm PVDF Transfer Packs, 1704156) at 25 V for 15 min. Membranes were blocked in blocking buffer (WB-BB, 5% w/v non-fat milk in 1x TBST buffer; Non-fat dry milk, RPI, M17200-500; 10x TBST, Genesee Scientific, 18-235B) for 1 h at room temperature and incubated overnight at 4 °C with indicated pAs diluted (1:1000 or 1:2000) in WB-BB. After washing 3x for 5 min each with 1x TBST, membranes were incubated with HRP-conjugated sAs (1:2000) diluted in WB-BB for 1 h at RT. Membranes were washed 3x with 1x TBST, subjected to chemiluminescent detection reagents (Biorad, 1705061 or 1705062) and imaged using a ChemiDoc imaging system. pAs used were as follows: rb-a-HSF1 (abcam, ab52757), rb-a-H3K27Ac (Invitrogen, 720096) and rb-a-HSF1-phospo S320 (abcam, ab76183). sAs used were as follows: goat-a-rb-HRP (CST, 7074S).

### Single-molecule RNA fluorescence in situ hybridization (smFISH)

Cells in 8 well chambered cover glass were fixed and washed as described in the IF section. Cells were then permeabilized with 70% ethanol at 4 °C for a minimum of 4 h. Following a brief wash, fixed cells were rehydrated for 5 min in smFISH wash buffer A (WBA10, 2:7:1 ratio of Wash Buffer A, nuclease-free water, and formamide; Wash buffer A, LGC Biosearch Technologies™, SMF-WA1-60; Nuclease Free Water, Invitrogen, 10-977-023; Formamide, Invitrogen, 15515-026). Probe hybridization was done by treating cells with smFISH hybridization solution (HybB10, Hybridization Buffer with 10% formamide; Hybridization Buffer, LGC Biosearch Technologies™, SMF-HB1-10) containing 100 nM custom-designed probes (LGC Biosearch Technologies™) and incubating them for 16 h at 37 °C, in a dark humidified chamber. Probe sequences are in Supplementary Table 1. After hybridization, cells were washed 1x with WBA10 at RT and washed again with WBA10 for 30 min at 37 °C, in a dark humidified chamber. Cells were washed again with concomitant nuclear staining using 20 ng/mL DAPI in WBA10 at 37 °C for 30 min, in a dark humidified chamber. Cells were then washed 1x with smFISH wash buffer B (WBB; LGC Biosearch Technologies™, SMF-WB1-20) for 5 min at RT, in the dark, and 1x with Tris-SSC buffer (TSSC, 10mM Tris.HCl, pH 7.6 with 2x SSC; 1M Tris.HCl, Invitrogen, 15567027; 20x SSC, Invitrogen, 15-557-044) for 5 min at RT, in the dark. Cells were mounted in smFISH antifade buffer (TSSC-TPP, as in PBS antifade, but with TSSC instead of PBS) and overlaid with low density mineral oil prior to imaging. For smFISH coupled with IF, smFISH was performed as above until the final wash step. Cells were re-refixed with 4% PFA for 10 min at RT in the dark and washed 3x with 1× PBS. IF was then carried out, starting from the permeabilization step and cells were finally mounted and imaged.

### 3-dimentional highly inclined laminated optical sheet (3D-HILO) microscopy

3D-HILO^49^ imaging was performed using a customized ONi (Oxford Nanoimager, Oxford Nanoimaging, UK) using a 100x / 1.49 NA oil immersion objective. The microscope is equipped with 405 nm, 473 nm, 532 nm and 647 nm lasers for excitation of fluorophores like DAPI, FITC/GFP, Cy3/JF549-HL and Cy5/JF646-HL/NIR Cell Mask respectively. All lasers are 1 W at the source with maximum achievable excitation power of 20 mW for 405 nm, 300 mW for 473 nm, 300 mW for 532 nm and 300 mW for 647 nm lasers at the objective. Custom multipass dichroics with reflectance at 309/40 nm, 408/20 nm, 532/14 nm, 640/14 nm and 850/20 nm, and transmission at 446/32 nm, 510/20 nm, 582/64 nm and 747/166 nm, was placed at 45°, below the objective to reflect lasers into the objective and transmit fluorescence. A series of 633 nm long pass filter and mirror were placed at 45° to split fluorescent light onto two 50 µm × 80 µm regions of a single CMOS camera. Multipass emission filter with band passes at 441/28 nm (e.g. for DAPI), 511/26 nm (e.g. for GFP) and 585/70 nm (e.g. for Cy3 or JF549-HL) was used at one channel and another with band passes at 704/78 nm (e.g. for Cy5 or JF646-HL) and 798/64 nm (e.g. for NIR Cell Mask) was used at the other channel. The microscope is capable of 4-color sequential imaging and 2-color simultaneous imaging at a lateral resolution of 30 nm and axial resolution of 100 nm with point-spread function modeling. The two channels used for sequential or simultaneous acquisition were first registered on the ONi via an automated version of previously described methods^50^. Registration was achieved by imaging 0.1 μm tetraspeck beads (Thermo-Fisher, T7279), at imaging conditions akin to those of cell samples. The registration matrix was then applied to the two channels images for accurate localization. For fixed cell imaging of dox. induced knock-ins and for IFs, 0.5 mW of 405 nm, 1 mW of 532 nm and 1 mW of 647 nm lasers were used to image DAPI, JF549-HL/Cy3 and Cy5/NIR Cell Mask respectively at 30 ms exposure. For smFISH of *HSP90AA1*, 0.5 mW of 405 nm, 5 mW of 532 nm and 5 mW of 647 nm lasers were used to image DAPI, Quasar 570 (on intron probes) and Quasar 670 (on exon probes) respectively at 30 ms exposure. For smFISH of *HSATIII*, 0.5 mW of 405 nm and 0.5 mW of 647 nm lasers were used to image DAPI and Cy5 respectively at 30 ms exposure.

### High-resolution imaging at high-throughput (HRIHT)

HRIHT was achieved using a customized microscope (Optical Biosystems, Inc) capable of performing synthetic-aperture optics assisted structured illumination microscopy (SAO-SIM)^51^ via a 20x / 0.45 NA air objective. The microscope is equipped with 365 nm, 473 nm, 532 nm and 647 nm lasers for excitation of fluorophores like DAPI, FITC/GFP, Cy3/JF549-HL and Cy5/JF646-HL/NIR Cell Mask respectively. Lasers have an excitation power of 500 mW for 473 nm, 2000 mW for 532 nm and 2000 mW for 647 nm at the source. Fluorescence was collected using bandpass filters – 460/36 nm (e.g. for DAPI), 520/40 nm (e.g. for FITC or GFP), 582/75 nm (e.g. Cy3 or JF549-HL), 708/75 nm (e.g. Cy5 or JF646-HL) and 810/90 nm (e.g. NIR Cell Mask) – across a CMOS camera. The field of view spans an area of 0.667 mm × 0.667 mm, enabling the simultaneous visualization of 50 – 100 cells, with a lateral resolution of 80 nm with point-spread function modeling. For fixed cell imaging of dox. induced knock-ins and for IFs, 200 mW of 532 nm and 200 mW of 647 nm lasers were used to image DAPI, JF549-HL/Cy3 and Cy5/NIR Cell Mask respectively at 50 ms exposure. Average image of patterned illumination, i.e. equivalent to epi-illumination on a wide-field microscope, was sufficient to visualize mesoscale condensates and these images were used for downstream analysis.

### Intra-condensate single-molecule tracking (SMT) and analysis

Dox. induced live knock-in cells in 8 well chambered cover glass were labelled with 0.2 nM JF549-HL and 200nM JF646-HL (gift from the Lavis Lab, JF646 Halotag ligand) diluted in phenol red free growth medium (phenol red free McCoy’s 5A modified medium with 10% FBS and 100 U/mL P-S; phenol red free McCoy’s 5A, Innovative Research Inc., SKU: IMCCOYS5A0714500ML) for 1 h, followed by 2x 15 min washes with phenol red free growth medium. After appropriate treatments in phenol red free growth medium, cells were imaged in live cell imaging medium (phenol red free McCoy’s 5A modified medium with 0.5% FBS and 100 U/mL P-S). Cells were then subjected to the appropriate stress treatments for indicated durations and used for imaging. For single-molecule tracking, chambered cover glass with live, labeled cells, was placed in a preheated (37°C typically or 42°C for HS), 5% CO_2_ controlled customed designed Tokai-Hit stage top incubator mounted in the ONi and imaged immediately. Cells were simultaneously excited with 45 mW of 532 nm laser and 0.5 mW of 640 nm laser at an exposure time of 30 ms.

SMT analysis was done as previously^52^, with minor modifications. Single JF549-HL particles within each cell were tracked in live-cell imaging videos using TrackMate^53^. A laplacian of gaussian (LoG) filter was used to aid localization of individual molecules and a LAP tracker with a maximum inter-frame displacement of 5 pixels (i.e. equivalent to the diameter of the particle), avoiding any gap filling was used for tracking. A bounding box (square or rectangle) enclosing the span of each condensate was visually chosen based on the JF646-HL signal. Boundaries of condensates were defined using a LoG filter and tracing contiguous intensity minima between condensates and adjacent background. Tracks within this bonding box were then used to calculate the mean squared displacement (MSD) and diffusion coefficients (D) using in-house MATLAB routines. All tracks were assumed to be Brownian. Only tracks longer than 15 frames (30 ms / frame) were used for diffusion coefficient calculations based on the equation: MSD = 4Dt^54^.

### Condensate and co-condensation analysis

For condensate analysis in 3D-HILO imaging (Supplementary Fig. 1), Z-stack images were first converted to 2D via maximum intensity projection. To obtain nuclei labels, the DAPI images were segmented with a fine-tuned Cellpose v3.1.0 model. HSF1 condensates were segmented with a distinct fine-tuned Cellpose v3.1.0 model. Briefly, a test dataset with hundreds of stressed nuclei and thousands of condensates were used to define condensates. This model was then applied to all data, which effectively demarcated condensates in stressed nuclei from the relatively diffuse localization under unstressed conditions. HSF1 condensate segmentation model served as an agnostic caller for condensates formed by other biomolecules (e.g. the transcription apparatus, i.e. H3K27ac, BRD4, MED12, RNAPII) as well. To standardize and parallelize condensate calling, the bounding box of each nucleus is padded with blank pixels to a standard size (256×26 pixels) and then rescaled using bi-linear interpolation (*s*=2) prior to segmentation. The number of condensates, apparent partition, nuclear mean fluorescent intensity (MFI), condensate area, and HSF1 condensate eccentricity is then derived from the image and nuclear and condensate labels. All analysis were performed with custom python scripts, predominantly supported by the NumPy v2.0.2 and skimage v0.25.0 libraries.

HSF1 condensate meta-images and metaplots were done as follows (Supplementary Fig. 2). For every HSF1 condensate label, an n-pixel square is extracted for both HSF1 signal and the corresponding co-imaging signal. For a given group of condensates, a mean image (meta-image) is created, and a background-subtracted (min. value) diagonal line scan is quantified. To compare the strength of a given enrichment meta-image or metaplot to cells without HSF1 condensates (e.g. mock-treated cells), random HSF1 condensates collected from a control condition (i.e. HS) are randomly placed onto the nuclei without condensates, where the number of condensates being placed is determined via sampling of a normal distribution derived from the actual control data. P-values are quantified for the metaplots by performing an unpaired T-test of the sum of co-imaging signal +/- 3 pixels from the center between conditions. For the HSF1 condensate area-matching analysis, condensates were filtered by area using Kolmogorov-Smirnov test thresholding and importance sampling to match HSF1 condensate area distributions across stresses to a control stress defined as the stress with the lowest mean HSF1 condensate area (i.e. ATO). Condensates passing this filtering were then used as input for metaplot creation as described above.

### smFISH image analysis

Nuclei segmentation was performed as previously described. To segment *HSP90AA1* intron foci, an iterative intensity thresholding method was used on each nucleus. Briefly, local maxima was iteratively identified at various thresholds that ranged from zero to maximum intensity of the image, at bin width of 100 intensity units. The optimal threshold for each nucleus was then determined with kneed v0.8.5. Foci were defined as contiguous non-zero pixels greater than the threshold. Putative transcription sites were defined as the top two segmented *HSP90AA1* intron foci with the greatest total fluorescent intensity (TFI) within each nucleus. Exon TFI was measured by overlaying the intron-derived mask for each nucleus on the corresponding *HSP90AA1* exon channel.

### C. elegans strain and maintenance

The following worm strain was sourced from the Caenorhabditis Genetics Center (CGC) : OG497 *unc-119(ed3);drSi13[hsf-1p::hsf-1::GFP::unc-54utr;Cb-unc-119+]*. Worms were maintained at 20℃ on nematode growth media (NGM) spotted with OP50 bacterial strain as a food source. Asynchronous populations of worms were washed off maintenance plates using 1–2 mL of M9 buffer and transferred into 1.7 mL Eppendorf tubes. To eliminate excess bacteria, worms were washed three times with fresh M9 solution (22.1 mM KH₂PO₄, 42.3 mM Na₂HPO₄, 85.6 mM NaCl, 1 mM MgSO₄), allowing them to settle by gravity between washes and carefully removing the supernatant each time. After the final wash, M9 was replaced with 1 mL hypochlorite solution (56 mL ddH₂O, 14.4 mL 5N NaOH, 7.2 mL 7.55% bleach). Tubes were shaken at 1200 rpm on a tabletop heat block set to 20 °C for 7–10 minutes, until no worm bodies were visible. Tubes with eggs only were centrifuged at 1600 rcf for 30 seconds, the hypochlorite solution was discarded, and the pellet was washed twice with fresh M9 solution. A final centrifugation was performed at 2900 rcf for 3 minutes, after which the M9 was removed. The egg pellet was resuspended in ∼100 μL M9 and transferred onto a fresh NGM plate seeded with OP50. Worms were cultured at 20 °C for approximately 72 hours to reach adulthood. Day 1 adults were used for subsequent experiments and imaging.

### C. elegans treatment, imaging, and analysis

Day 1 adult worms were individually transferred onto 2% agarose pads on glass slides containing 15–20 μL of either M9 solution, 1 mM SA (in M9), or 5 mM SA (in M9). Worms were incubated in the specified solution for 30 minutes, then immediately covered with a glass coverslip for imaging. Images were acquired as Z-stacks using a 40x/1.4 oil immersion objective on a Zeiss LSM 800 confocal laser scanning microscope. Granules were quantified in hypodermal and/or seam cell tail nuclei (4–15 nuclei per worm; 10–12 worms per condition). For the heat shock assay, synchronized worms were incubated at 37 °C for 30 minutes. After treatment, worms were individually placed on 2% agarose pads with 15–20 μL immobilization solution (25 mM tetramisole hydrochloride; Santa Cruz, sc-215963) and imaged immediately. For analysis, Z-stack images were first transformed with MIP on in-focus z-planes. Segmentation of nuclei and condensates was performed with a fine-tuned Cellpose models as previously described, except that the fine-tuning data utilized the HSF1::GFP worm images. Nuclear intensities and HSF1 apparent partition was calculated as described above.

### E. coli Spike-in DNA Preparation

A 5 ml liquid culture of DH5ɑ competent *E. coli* cells (Invitrogen) in LB buffer was made from glycerol stock and incubated overnight at 37°C, on shaker with rotation at 250 rpm. 1 ml of the liquid culture was spun down at 4500 xg for 5 min at RT in a swinging bucket centrifuge, resuspended in 600 uL of Lysis buffer (0.6% SDS and 0.12 mg/ml Proteinase K in 1X TE buffer) and incubated for 1.5h at 55C. An equal volume (600 ul) of phenol:chloroform:isoamyl alcohol was added to the sample, mixed by inversion, spun down at 20,000 xg for 5 min at RT in a tabletop centrifuge and the aqueous (top) phase collected in a new 1.5 ml tube. This step was repeated twice, once with an equal volume of phenol:chloroform:isoamyl alcohol and another with equal volume of chloroform. The final aqueous phase was incubated with 1 uL 10 mg/mL RNase A (Thermo Scientific) and 3 uL 20 mg/mL Proteinase K (GoldBio) for 1h at 37C, after which the phenol:chloroform:isoamyl alcohol and chloroform extraction steps were repeated. For DNA precipitation, 60 uL 5M NaCl and 1 mL isopropanol were added to the final aqueous phase and the sample incubated overnight at -20C. Sample was spun down at 20,000 xg for 15 min at RT in a tabletop centrifuge, the pellet was washed once in 1 ml ethanol, air dried for 5 min and then resuspended in 20 ul NF water. DNA concentration was measured on Qubit 4 fluorometer (Invitrogen) with High Sensitivity double stranded DNA reagents (Invitrogen) and the sample diluted to make 15 ul aliquots at 0.5 ng/uL of DNA concentration for fragmentation. Aliquots were sheared to 200 bp fragments in Covaris ME220 sonicator (Perkin Elmer) in microTUBEs-15 with AFA beads (Perkin Elmer), at the following conditions: 70s duration, 50W peak power, 30% duty factor, 50 cycles/burst and 15W average power. Final fragmented DNA size was assessed by Tapestation (Agilent), with HSD1000 reagents and tape (Agilent), and aliquots stored at -80°C.

### Cleavage under targets & release using nuclease (CUT&RUN)

CUT&RUN^55^ was performed according to the EpiCypher CUTANA CUT&RUN Protocol v1.6, with fresh whole cells. 11 ul of Concanavalin A (ConA) beads (EpiCypher) per CUT&RUN sample were transferred to a 1.5 ml tube and placed in a magnetic rack until the slurry cleared. Beads were washed twice in 100 μL per sample of Bead Activation buffer (20 mM HEPES-KOH, pH 7.9, 10 mM KCl, 1 mM CaCl_2_, 1 mM MnCl_2_), resuspended in 11 ul per sample of Bead Activation buffer, aliquoted into individual low-retention 1.5 ml tubes (10 ul per sample) and kept on ice until combined with samples. 500,000 freshly trypsinized cells were collected per CUT&RUN sample into low-retention 1.5 ml tubes, spun down at 600 xg for 3 min at RT in a swinging bucket centrifuge and washed once in 100 ul Wash Buffer (20 mM HEPES pH 7.5, 150 mM NaCl, 0.5 mM spermidine, 1x Roche cOmplete, Mini, EDTA-free Protease Inhibitor Cocktail). Cells were resuspended in 100 ul per sample of Wash Buffer, transferred to the PCR tubes containing activated ConA beads and incubated at RT for 10 min. The cells:beads were placed in a magnetic rack until the slurry cleared, resuspended in 50 ul Antibody buffer (2 mM EDTA in Digitonin buffer) with primary antibody (0.5 ug/sample of anti-HSF1 (Abcam #ab52757), anti-H3K36me3 (Cell Signaling #4909) or Rabbit IgG control (R&D Biosciences #AB-105-C)) and incubated for 1 h at RT, on nutator. Samples were washed twice in 200 ul Digitonin buffer (0.01% Digitonin in Wash Buffer). For MNase binding, cells:beads were resuspended in 50 ul Digitonin buffer with pAG-MNase (EpiCypher, 1:20) and incubated for 10 min at RT, on nutator, followed by two washes in 200 ul Digitonin buffer. Next, pAG-MNase was activated by incubation of cells:beads in 50 ul Digitonin buffer plus 1 ul of 100 mM CaCl2 for 2h at +4C, on nutator. MNase digestion was stopped by adding 33 ul Stop buffer (340 mM NaCl, 20 mM EDTA, 4 mM EGTA, 50 μg/mL RNase A, 50 μg/mL Glycogen) containing 0.5 ug *E. coli* spike-in DNA to each sample and incubation for 10 min at 37C in thermocycler. The supernatant containing the digested chromatin was transferred to a new PCR tube and incubated with 0.1% SDS and 14 ug Proteinase K (GoldBio) per sample for 10 min at 70C in thermocycler. Samples were frozen at -20C for extraction to be performed later.

For DNA extraction, samples were thawed on ice, transferred to a 1.5 ml low-retention tube and brought to a final volume of 250 ul per sample with 0.1X TE buffer. An equal volume (250 ul) of phenol:chloroform:isoamyl alcohol was immediately added per sample and mixed by inversion. Samples were spun down at 20,000 xg for 5 min at RT in a tabletop centrifuge and the aqueous (top) phase collected in a new low-retention 1.5 ml tube. Same volume of phenol:chloroform:isoamyl alcohol was added to samples, mixed by inversion, spun down at 20,000 xg for 5 min at RT in a tabletop centrifuge and the aqueous phase collected in a new low-retention 1.5 ml tube. For the final extraction step, same volume of chloroform was added to samples, mixed by inversion, spun down at 20,000 xg for 5 min at RT in a tabletop centrifuge and the aqueous phase collected in a new low-retention 1.5 ml tube. Extracted DNA was cleaned up with AMPure XP beads according to manufacturer’s instructions, at 2X ratio, and eluted in 12 ul NF water per sample. DNA concentration was measured on Qubit with High Sensitivity double stranded DNA reagents and samples were stored at -20C until library preparation. Libraries were made using SRSLY PicoPlus NGS Library Prep Kit (ClaretBio) according to manufacturer’s instructions and subjected 2×150 paired-end sequencing on the Illumina NovaSeq platform.

For HSF1 CUT&RUN analysis, sequenced reads were first aligned to the human genome with bwa-mem. To quantify fragment lengths, the BAMs were then converted to bedpe format with bedtools v2.26.0. A catalog of reads stemming from <= 120 bp fragments or >= 150 bp fragments was created then used to filter the using samtools v1.17. Each bam file was then converted to bigwig format using DeepTools BamCoverage v3.5.2 with bin size set to 1 and extending reads. The scale factor was set using the E. *coli* spike-in reads added to each sample. To calculate the spike-in factor for each sample, first, the fastqs were aligned to the E. *coli* genome and the total number of reads were quantified with featureCounts v2.0.6. Then the spike-in factor was calculated as 1 / (E. *coli* reads/ Total reads). To create the metaplots, first, HSF1 peaks were identified using SEACR v1.3 with the following parameters, cutoff-0.1, norm = “non”, and mode = “stringent”. To identify HSEs within peaks, a comprehensive list of HSF1 SEACR peaks was compiled, then their sequences were used as input to FIMO implemented by memesuite-lite v0.2.0 with parameters, p=1e-5. Metaplots were created using pyBigWig v0.3.24 to extract signal around given HSEs for each sample and then calculating the mean across replicates. To group HSEs by regulatory pattern, the log2FC of mean signal at each HSE +/- 50 bp was calculated for each stress sample compared to the mock-treated signal. HSEs with > 0 log2FC were defined as upregulated. For plotting, differences imparted in signal by the spike-in were accounted for by subtracting the background from each HSE loci (+/- 1kb) defined as the minimum value in each region.

### Nascent RNA sequencing (Bru-Seq)

For Bru-seq, cells were mock treated or subjected to HS or SA treatment (as above). At the final 30 min of mock treatment or stress, cells were washed once with fresh medium before bromouridine (BrU) treatment. BrU solution was diluted to a final concentration of 2 mM in growth medium and added to cells for 30 min. Nascent transcript libraries for Bru-seq were prepared and sequenced as described^56^. Sequenced reads were first aligned to the human genome with STAR v2.7.10b. To create the metaplot, each bam file was converted to bigwig format using deepTools BamCoverage. Then, pyBigWig was used to extract the signal at the given genes’ TSS +/- 2 kb and the mean at each position was calculated. For differential gene expression, limma from Bioconductor v3.18 was used. Euler diagrams were created with the eulerr python library.

### Statistical analysis

All experiments were done at least three times. Statistical analyses were performed using Graphpad Prism v10, R package 4.3 and Python v3.8.8. Pairwise comparisons were done using student’s t-tests, wherein p < 0.05 was considered significant. No data were excluded from analyses.

### Data availability

All source data in this study are available will be made freely available upon publication in a peer-reviewed journal. Sequencing data have been deposited in the Gene Expression Omnibus (accession number GSE319297, and GSE319125) and are currently private, but access can be provided upon request. All source codes can be made available upon request.

## ACKNOWLEDGEMENTS

We thank Lakshmi Dommeti, Ramya Ravishankar and Nivea Vydiswaran for early technical assistance. S.R. was supported by NIH R35 GM133434 and R35 GM156411. J.N.D. was supported by NIH T32 GM145470. BrU-Seq was supported by NCI 2P30CA046592-34. T.H. was supported by China Postdoctoral Science Foundation (Grant No. 2025M781888). This work was supported in part by the Urology Care Foundation Physician Scientist Residency Training Award program and Dornier MedTech to J.E.B. This study was supported by NIH R35 GM155432, NIH/NIDDK grant U54DK137314 pilot award, Urology Catalyst Award, and funds from the Department of Urology, Rogel Cancer Center and the Michigan Center for Translational Pathology to S.P.

## AUTHOR CONTRIBUTIONS

S.P. conceived and designed the study. J.E.B., C.K.D. and A.S. performed imaging and biochemical assays. J.N.D. analyzed all imaging and omics data. T.H. and L.X. constructed plasmids and created cell lines. G.B.V., S.R. and M.L. performed sequencing assays. B.A. and M.T. performed *C. Elegans* assays. S.P. and J.N.D. wrote the manuscript and all authors provided feedback.

## COMPETING INTERESTS

No competing interests were declared from the authors.

## MATERIALS & CORRESPONDENCE

For all material requests please contact.

## Notes

### Competing Interest Statement

The authors have declared no competing interest.

