## Supplementary Figures for "Divergent condensates tune transcriptional responses during stress"

#### SUPPLEMENTARY MATERIALS

##### Supplementary Figure 1

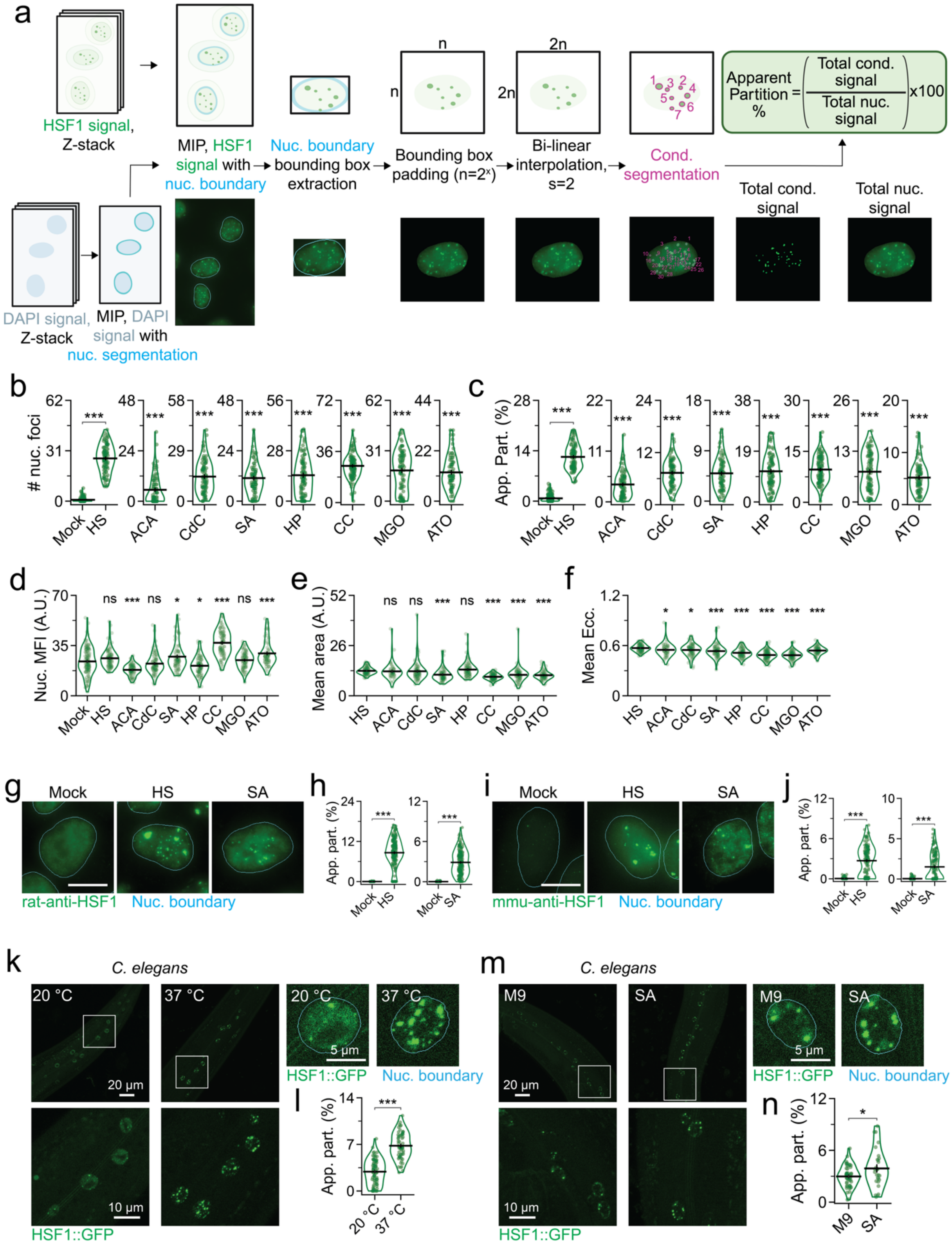

**Supplementary Figure 1. HSF1 condensate formation is an evolutionarily conserved feature of stress response.**

(a) Schematic of HSF1 foci analysis.

(b) Quantified number of HSF1 foci per nucleus (nuc.). ***=$p<0.0005$, by two-sided unpaired T-test, as compared to Mock. Line represents mean and error bars represent s.e.m. Mean is represented in Fig. 1b.

(c) Quantified HSF1 apparent partition (App. Part.) per nuc. ***=$p<0.0005$, by two-sided unpaired T-test, as compared to Mock. Line represents mean and error bars represent s.e.m. Mean is represented in Fig. 1b.

(d) Quantified HSF1 nuc. mean fluorescent intensity (MFI). *=$p<0.05$, ***=$p<0.0005$, ns = not significant, by two-sided unpaired T-test, as compared to Mock. Line represents mean and error bars represent s.e.m. Mean is represented in Fig. 1b.

(e) Quantified mean HSF1 foci area per nuc. *=$p<0.05$, ***=$p<0.0005$, ns = not significant, by two-sided unpaired T-test, as compared to HS. Line represents mean and error bars represent s.e.m. Mean is represented in Fig. 1b.

(f) Quantified mean HSF1 foci eccentricity (Ecc.) per nuc. *=$p<0.05$, ***=$p<0.0005$, by two-sided unpaired T-test. Line represents mean and error bars represent s.e.m. Mean is represented in Fig. 1b.

(g) Representative pseudo-colored IF images of U2OS treated that were mock treated or subjected to HS, or SA, with rat-anti-HSF1 (green). Scale bar, 10 μm.

(h) Quantified HSF1 foci number and App. Part. per nuc. from (g). ***=$p<0.0005$, by two-sided unpaired T-test ($n \geq 147$ cells, 2495 condensates, per condition). Line represents mean and error bars represent s.e.m.

(i) Representative pseudo-colored IF images of U2OS treated that were mock treated or subjected to HS, or SA, and stained with mmu-anti-HSF1 (green). Scale bar, 10 μm.

(j) Quantified HSF1 foci number and App. Part. per nuc. from (i). ***=$p<0.0005$, by two-sided unpaired T-test ($n \geq 145$ cells, 1251 condensates). Line represents mean and error bars represent s.e.m.

(k) Representative pseudo-colored fluorescence images of HSF1::GFP (green) in C. *elegans* at 20 °C (Mock) or 37 °C (HS, 1 h).

(l) Quantified HSF1 App. Part. from (k). ***=$p<0.0005$, by two-sided unpaired T-test ($n \geq 62$ cells, 331 condensates, per condition). Line represents mean and error bars represent s.e.m.

(m) Representative pseudo-colored fluorescence images of HSF1::GFP (green) in C. *elegans* that were mock treated (M9 buffer) or subjected to SA (1 mM, 30 m).

(n) Quantified HSF1 App. Part. from (m). *=$p<0.05$, by two-sided unpaired T-test ($n \geq 29$ cells, 112 condensates, per condition). Line represents mean and error bars represent s.e.m.

**Supplementary Figure 2**

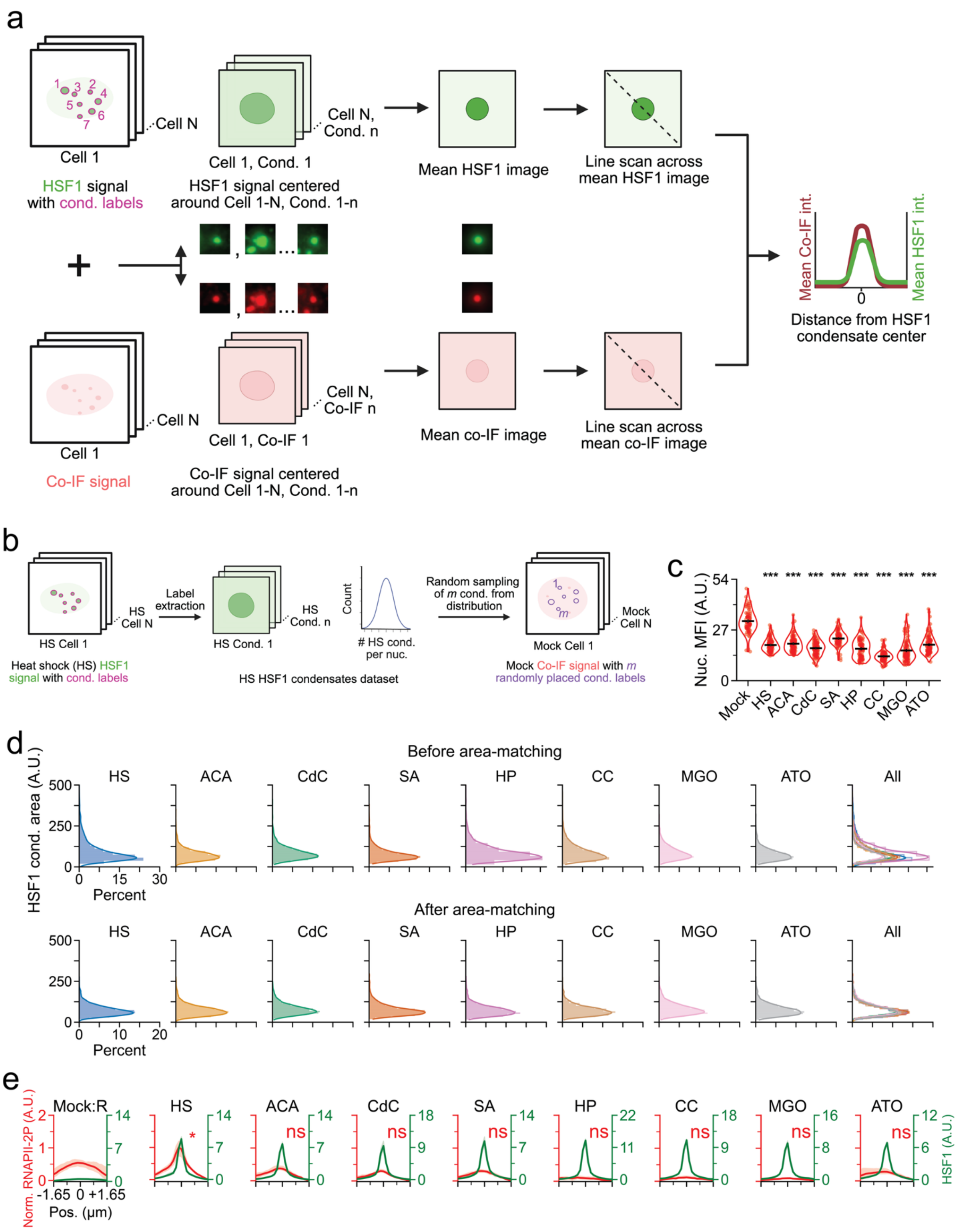

**Supplementary Figure 2. RNAPII-2P levels or extent of HSF1 condensation are not associated with stress-type dependent divergence in RNAPII-2P co-enrichment.**

(a) Schematic of HSF1 condensate metaplot analysis.

(b) Schematic of HSF1 condensate randomization.

(c) Quantification of nuc. RNAPII-2P MFI Fig. 1c. ***=$p<0.0005$, by two-sided unpaired T-test. Line represents mean and error bars represent s.e.m.

(d) Histogram of HSF1 condensate areas from Fig. 1c before and after area-matching.

(e) Line scan metaplots of RNAPII-2P across segmented, area-matched HSF1 condensates from Fig. 1c. *=$p<0.05$, ns = not significant, by two-sided unpaired T-test.

**Supplementary Figure 3**

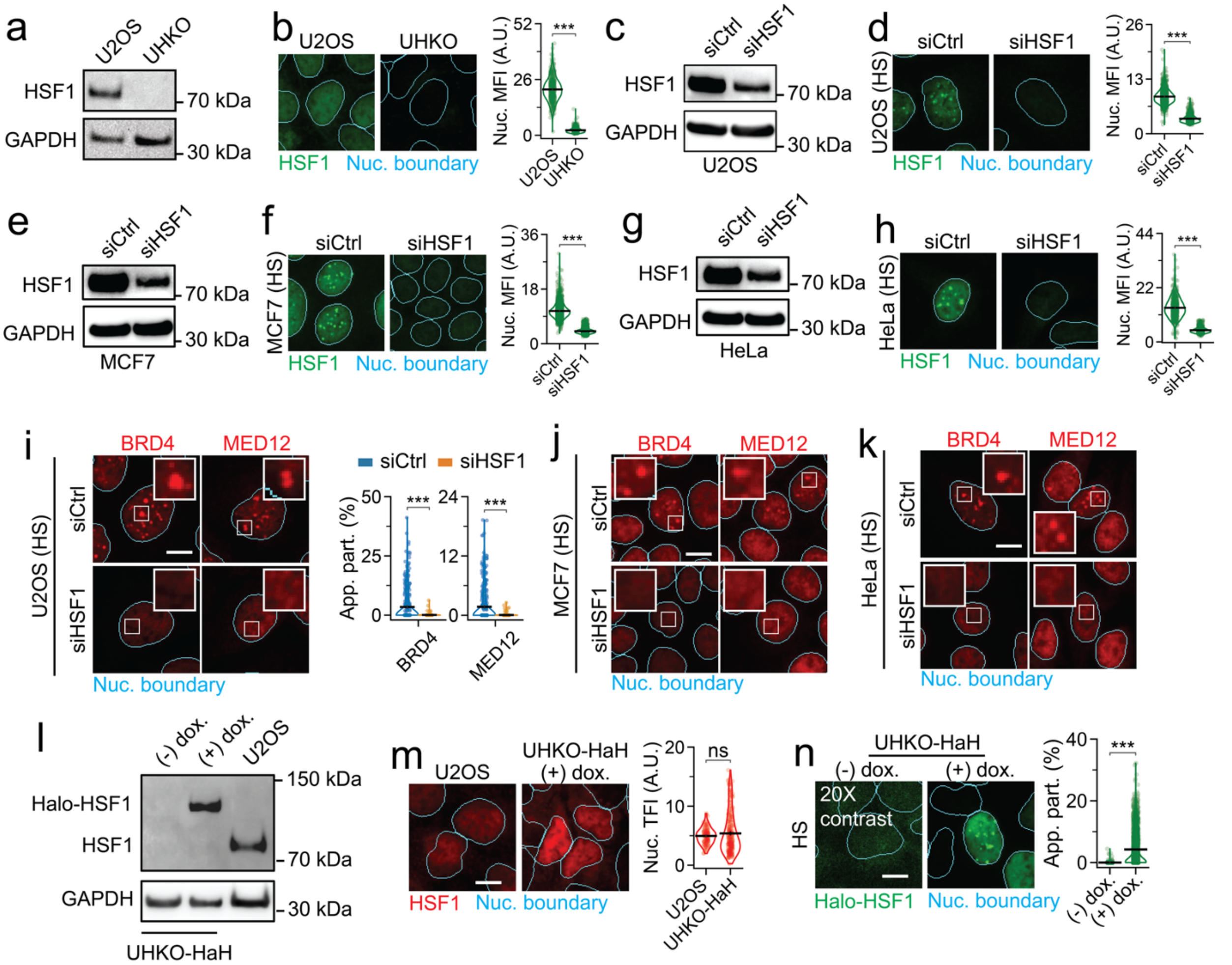


**Supplementary Figure 3. Benchmarking knockout, knockdown and inducible expression of HSF1.**

(a) Immunoblotting of HSF1 and GAPDH from U2OS and UHKO cells.

(b) Representative pseudo-colored IF images of U2OS and UHKO cells stained for HSF1 (green). Quantification of HSF1 Nuc. MFI is also shown. ***=p<0.0005, by two-sided unpaired T-test (n ≥ 912 cells, per condition). Line represents mean and error bars represent s.e.m.

(c) Immunoblotting of HSF1 and GAPDH from siCtrl or siHSF1 treated U2OS cells.

(d) Representative pseudo-colored IF images of U2OS cells treated with siCtrl or siHSF1, subjected to HS, and stained for HSF1 (green). Quantification of HSF1 nuc. MFI is also shown. ***=p<0.0005, by two-sided unpaired T-test (n ≥ 2457 cells, per condition). Line represents mean and error bars represent s.e.m.

(e) Immunoblotting of HSF1 and GAPDH from siCtrl or siHSF1 treated MCF7 cells.

(f) Representative pseudo-colored IF images of MCF7 cells treated with siCtrl or siHSF1, subjected to HS, and stained for HSF1 (green). Quantification of HSF1 nuc. MFI is also shown. ***=p<0.0005, by two-sided unpaired T-test (n ≥ 6135 cells, per condition). Line represents mean and error bars represent s.e.m.

(g) Immunoblotting of HSF1 and GAPDH from siCtrl or siHSF1 treated HeLa cells.

(h) Representative pseudo-colored IF images of HeLa cells treated with siCtrl or siHSF1, subjected to HS, and stained for HSF1 (green). Quantification of HSF1 nuc. MFI is also shown. ***=p<0.0005, by two-sided unpaired T-test (n ≥ 1415 cells, per condition). Line represents mean and error bars represent s.e.m.

(i) Representative pseudo-colored IF images of U2OS cells treated with siCtrl or siHSF1, subjected to HS, and stained for BRD4 or MED12 (red). Scale bar, 10 µm. Zoom ins are 5 µm x 5 µm. Quantification of BRD4 and MED12 App. Part. is also shown. ***=p<0.0005, by two-sided unpaired T-test (n ≥ 662 cells, per condition). Line represents mean and error bars represent s.e.m.

(j) Representative pseudo-colored IF images of MCF7 cells treated with siCtrl or siHSF1, subjected to HS, and stained for BRD4 or MED12 (red), from Fig. 2e. Scale bar, 10 µm. Zoom ins are 5 µm x 5 µm.

(k) Representative pseudo-colored IF images of HeLa cells treated with siCtrl or siHSF1, subjected to HS, and stained for BRD4 or MED12 (red), from Fig. 2f. Scale bar, 10 μm. Zoom ins are 5 μm x 5 μm.

(l) Immunoblotting of HSF1 and GAPDH from U2OS cells and UHKO-HaH cells treated with or without doxycycline (dox.).

(m) Representative pseudo-colored IF images of U2OS or dox.-induced UHKO-HaH cells stained for HSF1 (red). Quantification of HSF1 nuc. total fluorescent intensity (TFI) is also shown. ns = not significant, by two-sided unpaired T-test ($n \geq 127$ cells, per condition). Line represents mean and error bars represent s.e.m.

(n) Representative pseudo-colored fluorescent images of UHKO-HaH cells subjected to HS, treated with or without dox. and stained for Halo-HSF1 (JF549-HL, green). Contrast of (-)dox. image is 20x improved to show basal, diffuse, background signal. Quantification of HSF1 App. Part. is also shown. ***=$p<0.0005$, by two-sided unpaired T-test ($n \geq 1644$ cells, per condition). Line represents mean and error bars represent s.e.m.

**Supplementary Figure 4**

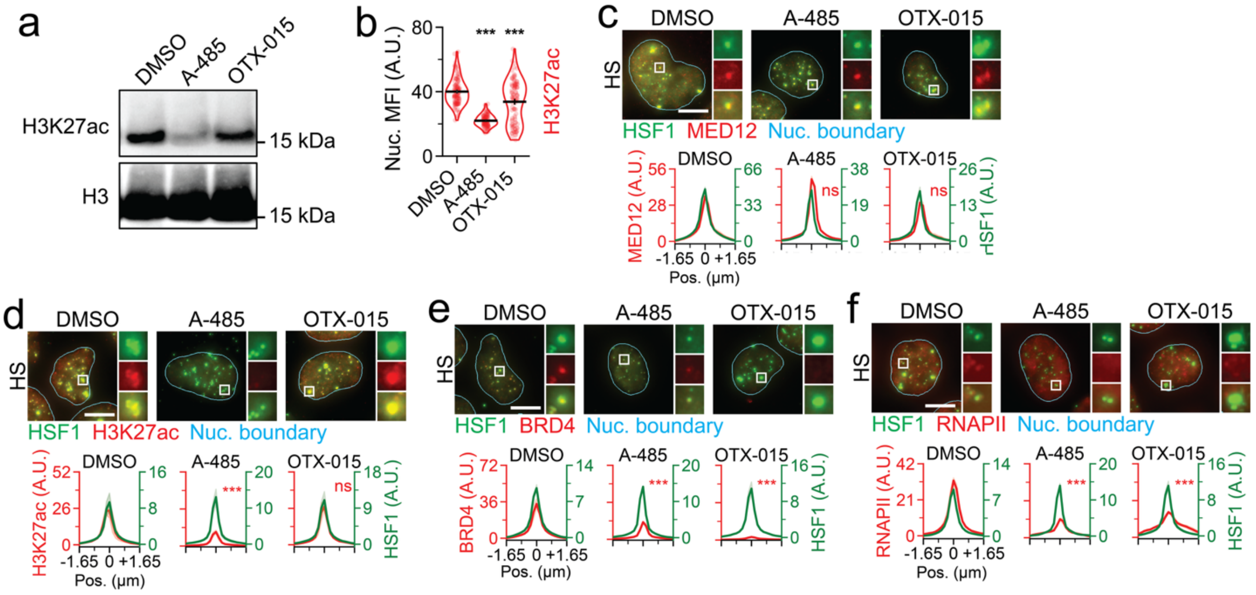


**Supplementary Figure 4. Epigenetic modulators affect HSF1 condensate function, but not formation.**

(a) Immunoblotting of H3K27ac and H3 of U2OS cells treated with DMSO, A-485, or OTX-015 and subjected to HS.

(b) Quantification of nuc. H3K27ac MFI in U2OS cells treated with DMSO, A-485, or OTX-015 and subjected to HS. ***=p<0.0005, by two-sided unpaired T-test (n ≥ 100 cells, per condition). Line represents mean and error bars represent s.e.m.

(c) Representative pseudo-colored IF images of U2OS cells treated with DMSO, A-485, or OTX-015, subjected to HS, and stained for HSF1 (green) and MED12 (red). Line scan metaplots of MED12 signal across segmented HSF1 condensates is also shown. ns = not significant, by two-sided unpaired T-test (n ≥ 101 cells, per condition). Meta-images are in Fig. 2k.

(d) Representative pseudo-colored IF images of U2OS cells treated with DMSO, A-485, or OTX-015, subjected to HS, and stained for HSF1 (green) and H3K27ac (red); Line scan metaplots of

H3K27ac signal across segmented HSF1 condensates is also shown. ***=p<0.0005, ns = not significant, by two-sided unpaired T-test (n ≥ 100 cells, per condition). Meta-images are in Fig. 2k.

(e) Representative pseudo-colored IF images of U2OS cells treated with DMSO, A-485, or OTX-015, subjected to HS, and stained for HSF1 (green) and BRD4 (red); Line scan metaplots of BRD4 signal across segmented HSF1 condensates is also shown. ***=p<0.0005, by two-sided unpaired T-test (n ≥ 90 cells, per condition). Meta-images are in Fig. 2k.

(f) Representative pseudo-colored IF images of U2OS cells treated with DMSO, A-485, or OTX-015, subjected to HS, and stained for HSF1 (green) and RNAPII (red); Line scan metaplots of RNAPII signal across segmented HSF1 condensates is also shown. ***=p<0.0005, by two-sided unpaired T-test (n ≥ 100 cells, per condition). Meta-images are in Fig. 2k.

#### Supplementary Figure 5

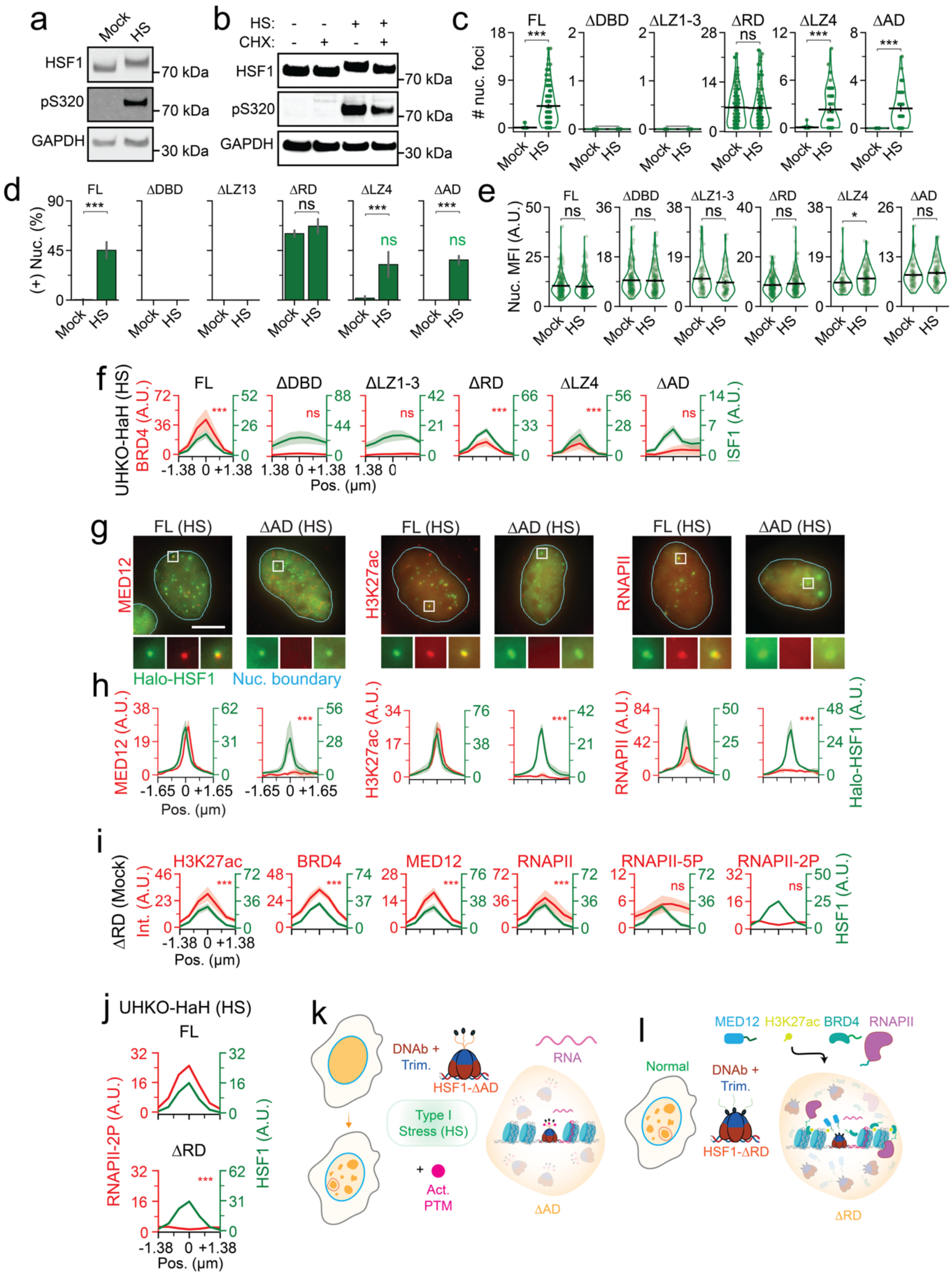

**Supplementary Figure 5. PTMs, DNA binding and trimerization drive condensate formation, but disordered regions inhibit condensation in unstressed situations, whilst enabling transcriptional function during stress.**

(a) Immunoblotting of HSF1, HSF1-pS320 (pS320) and GAPDH of U2OS cells that were mock treated or subjected to HS.

(b) Immunoblotting of HSF1, HSF1-pS320 (pS320) and GAPDH of U2OS cells treated with cycloheximide (CHX) and mock treated or subjected to HS.

(c) Quantification of number Halo-HSF1 foci per nuc. in UHKO-HaH cells from Fig. 3d. ***=p<0.0005, ns = not significant, by two-sided unpaired T-test. Line represents mean and error bars represent s.e.m.

(d) Quantification of the percentage of Halo-HSF1 foci positive (+ Nuc %) from Fig 3d. ***=p<0.0005, ns = not significant, by two-sided unpaired T-test. ns (green) = not significant, by two-sided unpaired T-test, when compared to HS. Error bars represent s.e.m.

(e) Quantification of Halo-HSF1 nuc. MFI from UHKO-HaH cells in Fig 3d. *=p<0.05, ns = not significant, by two-sided unpaired T-test. Line represents mean and error bars represent s.e.m.

(f) Line scan metaplots of MED12, H3K27ac or BRD4 signal across segmented Halo-HSF1 condensates in cells from Fig 3f. ΔDBD, ΔLZ1-3 plots are created with randomized condensates as previously described. ***=p<0.0005, ns = not significant, by two-sided unpaired T-test, when compared to randomizations.

(g) Representative pseudo-colored images of HS subjected FL and ΔAD cells stained with Halo-HSF1 (JF549-HL, green) and MED12, H3K27ac, or RNAPII (red). Meta-images are in Fig. 3g. Scale bar, 10 μm. Zoom-ins and meta-images are 2.7 μm x 2.7 μm.

(h) Line scan metaplots of BRD4 signal across segmented Halo-HSF1 condensates in cells from (g). ***=p<0.0005, by two-sided unpaired T-test, when compared to FL.

(i) Line scan metaplots of H327ac, BRD4, MED12, RNAPII, RNAPII-5P, and RNAPII-2P signal across segmented Halo-HSF1 condensates in mock-treated ΔRD cells from Fig. 3g. ***=p<0.0005, ns = not significant, by two-sided unpaired T-test, when compared to randomizations.

(j) Line scan metaplot of RNAPII-2P signal across segmented Halo-HSF1 condensates in HS-treated FL and ΔRD cells from Fig. 3h. ***=p<0.0005, by two-sided unpaired T-test, when compared to FL.

(k) Model representing lack of transcription machinery in HSF1-ΔAD condensates during HS.

(l) Model representing transcriptionally incompetent, stalled HSF1-ΔRD hubs that form even in the absence of stress.

### Supplementary Figure 6

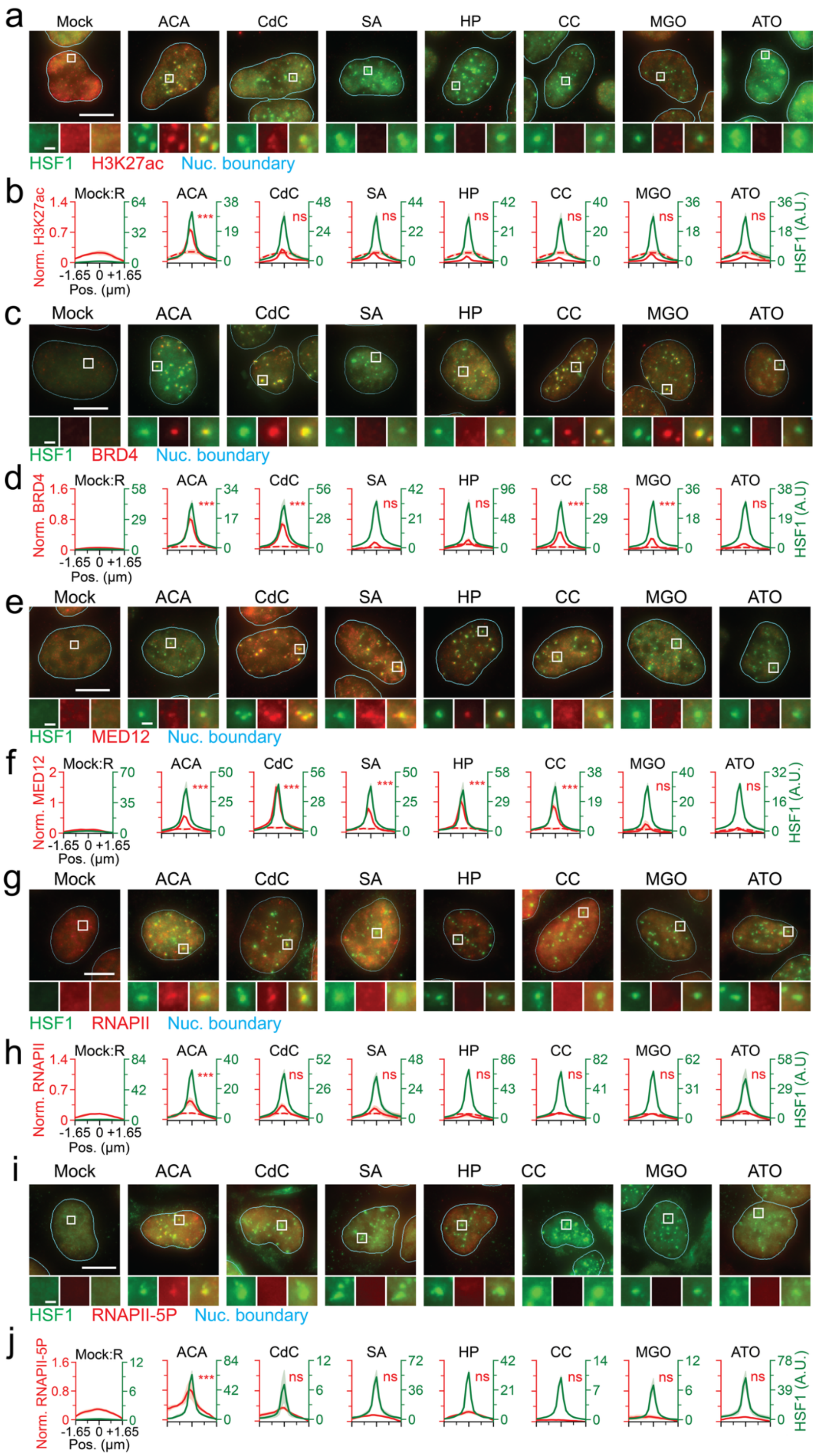

**Supplementary Figure 6. Full complement of factors required for active hubs are missing in HSF1 condensates that form during other, non-HS stresses.**

(a) Representative pseudo-colored IF images of U2OS cells treated with various stresses and stained for HSF1 (green) and H3K27ac (red). Meta-images are in Fig. 4a. Scale bar, 10 μm. Zoom-ins and meta-images are 2.7 μm x 2.7 μm.

(b) Normalized line scan metaplots of H3K27ac signal across segmented HSF1 condensates in cells from (a). ***=p<0.0005, ns = not significant, by two-sided unpaired T-test, when compared to randomizations in Mock.

(c) Representative pseudo-colored IF images of U2OS cells treated with various stresses and stained for HSF1 (green) and BRD4 (red). Meta-images are in Fig. 4a. Scale bar, 10 μm. Zoom-ins and meta-images are 2.7 μm x 2.7 μm.

(d) Normalized line scan metaplots of BRD4 signal across segmented HSF1 condensates in cells from (a). ***=p<0.0005, ns = not significant, by two-sided unpaired T-test, when compared to randomizations in Mock.

(e) Representative pseudo-colored IF images of U2OS cells treated with various stresses and stained for HSF1 (green) and MED12 (red). Meta-images are in Fig. 4a. Scale bar, 10 μm. Zoom-ins and meta-images are 2.7 μm x 2.7 μm.

(f) Normalized line scan metaplots of MED12 signal across segmented HSF1 condensates in cells from (a). ***=p<0.0005, ns = not significant, by two-sided unpaired T-test, when compared to randomizations in Mock.

(g) Representative pseudo-colored IF images of U2OS cells treated with various stresses and stained for HSF1 (green) and RNAPII (red). Meta-images are in Fig. 4a. Scale bar, 10 μm. Zoom-ins and meta-images are 2.7 μm x 2.7 μm.

(h) Normalized line scan metaplots of RNAPII signal across segmented HSF1 condensates in cells from (a). ***=p<0.0005, ns = not significant, by two-sided unpaired T-test, when compared to randomizations in Mock.

(i) Representative pseudo-colored IF images of U2OS cells treated with various stresses and stained for HSF1 (green) and RNAPII-5P (red). Meta-images are in Fig. 4a. Scale bar, 10 μm. Zoom-ins and meta-images are 2.7 μm x 2.7 μm.

(j) Normalized line scan metaplots of RNAPII-5P signal across segmented HSF1 condensates in cells from (a). ***=p<0.0005, ns = not significant, by two-sided unpaired T-test, when compared to randomizations in Mock.

**Supplementary Figure 7**

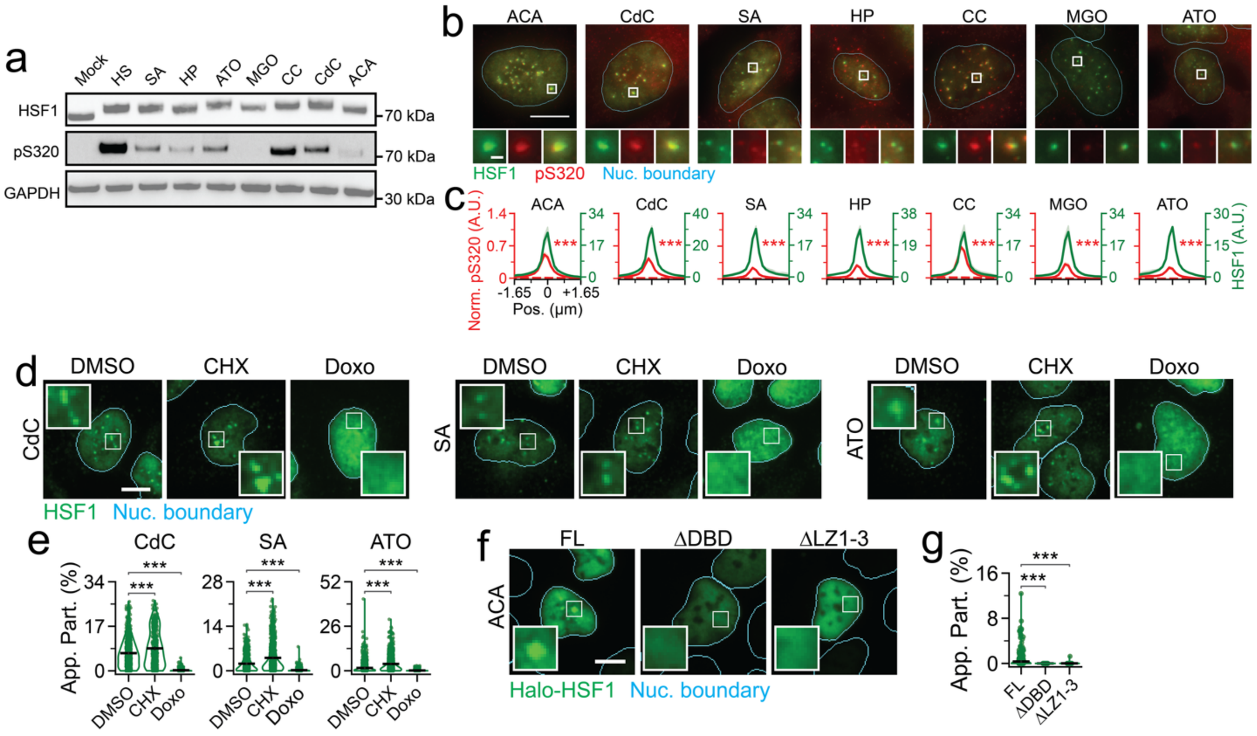


**Supplementary Figure 7. Transcriptionally incompetent HSF1 condensates in non-HS stresses require DNA binding and trimerization but form independent of PTMs.**

(a) Immunoblotting of HSF1, HSF1-pS320 (pS320) and GAPDH in U2OS cells subjected to various stresses.

(b) Representative pseudo-colored IF images of U2OS cells treated with various environmental and chemotherapeutic stresses and stained for HSF1 (green) and HSF1-pS320 (pS320, red). Scale bar, 10 µm. Zoom-ins and meta-images are 2.7 µm x 2.7 µm.

(c) Normalized line scan metaplots of HSF1-pS320 signal across segmented HSF1 condensates in (b). ***=$p<0.0005$, by two-sided unpaired T-test, when compared to randomizations in Mock.

(d) Representative pseudo-colored IF images of U2OS cells treated with DMSO, CHX, or Doxo and subjected to CdC, SA or ATO stress. stained for HSF1 (green). Nuc. Boundary (cyan) is derived from DAPI staining. Scale bar, 10 µm. Zoom ins are 5 µm x 5 µm.

(e) Quantification of HSF1 App. Part. in cells from (d). ***=$p<0.0005$, by two-sided unpaired T-test ($n \geq 409$ cells, per condition). Line represents mean and error bars represent s.e.m.

(f) Representative pseudo-colored fluorescent images of UHKO-HaH (FL, ΔDBD, and ΔLZ1-3) cells treated with ACA and stained for Halo-HSF1 (JF549-HL, green). Nuc. Boundary (cyan) is derived from DAPI staining. Scale bar, 10 μm. Zoom ins are 5 μm x 5 μm.

(g) Quantification of HSF1 App. Part. in cells from (f). ***=$p<0.0005$, by two-sided unpaired T-test ($n \geq 120$ cells, per condition). Line represents mean and error bars represent s.e.m.

**Supplementary Figure 8**

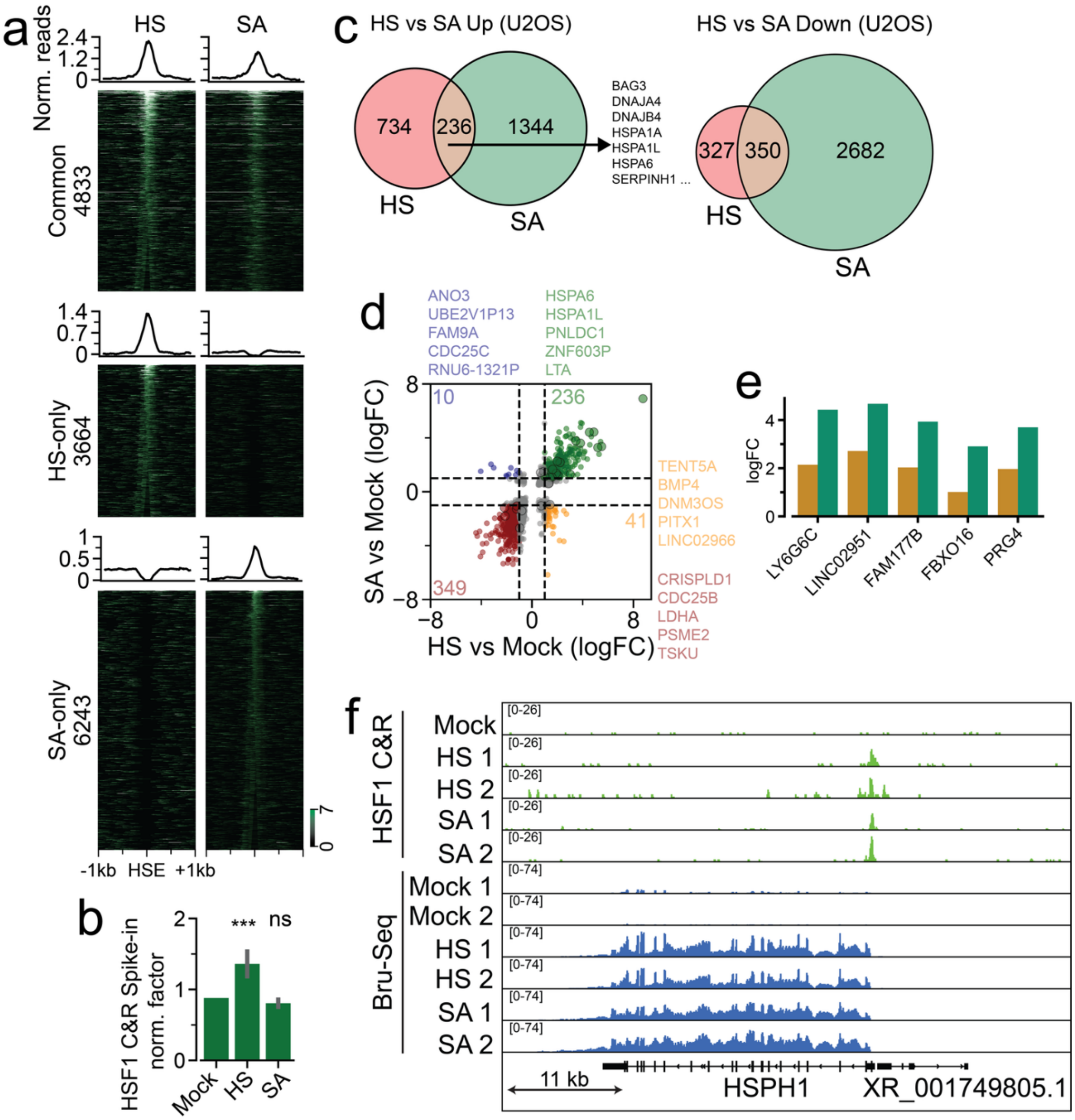


**Supplementary Figure 8. HSF1 cistrome and the transcriptome reorganize into context-agnostic and stress type-specific programs during HS and SA stress.**

(a) Spike-in normalization factor of HSF1 CUT&RUN data. ***=p<0.0005, ns = not significant, by two-sided unpaired T-test. Error bars represent s.e.m.

(b) Metaplots and heatmaps of ≤ 120 bp fragments HSF1 CUT&RUN signal grouped by pattern of upregulation, when compared to unstressed controls.

(c) Euler diagram of up- and down-regulated genes in Bru-Seq data of HS and SA-treated U2OS cells.

(d) Correlation scatter plot of differentially expressed genes in HS and SA as compared to Mock. Top right, top left, bottom right and bottom left quadrants of the plot represent transcripts that are significantly (adjusted p-val < 0.05) upregulated in both HS and SA ($log_2FC \geq 1$, green), uniquely upregulated in SA ($log_2FC \geq 1$ in SA and $log_2FC \leq -1$ in HS, blue), uniquely upregulated in HS ($log_2FC \geq 1$ in HS and $log_2FC \leq -1$ in SA, orange) and commonly downregulated in both HS and SA ($log_2FC \leq -1$, red), respectively. Color coded numbers indicate number of transcripts that pass cut-offs. Grey dots represent significantly regulated genes that did not pass the $log_2FC$ cut-offs. Top five genes pertaining to each quadrant are also listed.

(e) Representative list of 5 genes commonly upregulated in HS and SA, with extent of upregulation being higher in SA over HS.

(f) Representative tracks of HSF1 CUT&RUN and Bru-Seq data at *HSPH1* gene locus representing similar extents of HSF1 occupancy and *HSPH1* gene induction.

**Supplementary Video 1. Single-molecule tracking of HSF1 within and outside of condensates during HS.** Representative video of dox. induced and HS subjected UHKO-HaH cells with sparsely (JF549-HL, green) and densely (JF646-HL, red) labeled Halo-HSF1. Log-filtered rendition is represented to aid visualization of molecules and condensates. Tracks are color-coded by increasing displacement across a blue-red spectrum. Individual channels and merged versions are shown.

**Supplementary Table 1.** Sequences of CRISPR-Cas9 guides (DNAs) used to generate UHKO cells and smFISH probes.
